# Patterning the embryonic pulmonary mesenchyme

**DOI:** 10.1101/2020.08.20.259101

**Authors:** Katharine Goodwin, Jacob M. Jaslove, Hirotaka Tao, Min Zhu, Sevan Hopyan, Celeste M. Nelson

## Abstract

Smooth muscle guides morphogenesis of several epithelia during organogenesis, including the mammalian airways. However, it remains unclear how airway smooth muscle differentiation is spatiotemporally patterned and whether it originates from distinct mesenchymal progenitors. Using single-cell RNA-sequencing of embryonic mouse lungs, we show that the pulmonary mesenchyme contains a continuum of cell identities, but no distinct progenitors. Transcriptional variability correlates with sub-epithelial and sub-mesothelial mesenchymal compartments that are regulated by Wnt signaling. Live-imaging and tension-sensors reveal compartment-specific migratory behaviors and cortical forces, and show that sub-epithelial mesenchyme contributes to airway smooth muscle. Cytoskeletal and Wnt signaling pathways are activated early in reconstructed differentiation trajectories. Consistently, Wnt activation stimulates the earliest stages of smooth muscle differentiation and induces local accumulation of mesenchymal F-actin, which influences epithelial morphology. Our single-cell approach uncovers the principles of pulmonary mesenchymal patterning during branching morphogenesis and identifies a morphogenetically active mesenchymal layer that sculpts the airway epithelium.

## Introduction

Smooth muscle and similar contractile cell types influence the morphogenesis of epithelial tissues^1–3^. Branching of the mammalian airway epithelium has long been thought to involve smooth muscle differentiation from the surrounding pulmonary mesenchyme^4–8^. Accordingly, ex vivo culture revealed that patterned airway smooth muscle differentiation specifies cleft sites, promotes epithelial bifurcation, and shapes the domain branches that establish the underlying architecture of the murine lung^9,10^. It is therefore surprising that the smooth muscle transcription factor myocardin (*Myocd*) is dispensable for lung branching morphogenesis^11^ and that mice lacking α-smooth muscle actin (αSMA or *Acta2*) are viable and only develop deleterious phenotypes in smooth muscle and similar contractile tissues under conditions of high mechanical stress^12,13^. These studies suggest that smooth muscle differentiation and transcriptional identity are more nuanced that previously assumed, and highlight the importance of uncovering the factors that define a smooth muscle cell and control its differentiation.

Lung development is intricately regulated by signaling between the epithelium, mesothelium, and mesenchyme. Shh signaling from the epithelium is required for airway smooth muscle differentiation^14^, while Fgf9 signaling from the mesothelium prevents the differentiation of this tissue^15^. The physical layout of the airway smooth muscle and the differing diffusible signals from the epithelium and mesothelium could potentially establish separate mesenchymal compartments. In particular, it appears that epithelial and mesothelial signals directly influence the behavior of the sub-epithelial and sub-mesothelial mesenchyme, respectively^16,17^.

Cells from different locations within the pulmonary mesenchyme possess different capacities for differentiation: mesenchyme that is transplanted from stalk to tip regions fails to migrate towards the epithelium or differentiate into airway smooth muscle unless it is exposed to Wnt1^18^. Wnt signaling may therefore be involved in a compartment-specific manner in specifying airway smooth muscle progenitor identity and behavior. Consistently, *Wnt2a* is expressed in the distal mesenchyme^19^ and the expression of *Axin2* is strongest in the mesenchyme around branch tips^20^. Deleting *Ctnnb1* (β-catenin) from the mesenchyme delays branching morphogenesis and inhibits mesenchymal growth and survival^17,21,22^. Expressing stabilized β-catenin leads to disorganized airway smooth muscle wrapping^18^, and Wnt2 promotes the expression of myogenic transcription factors to support robust smooth muscle differentiation^23^.

Lineage-tracing studies based on single marker genes have not identified a distinct airway smooth muscle progenitor^24,25^. Additionally, the signals required for patterned smooth muscle differentiation remain unclear. Multiple pathways have been implicated but studying each one individually will not reveal how they interact to achieve spatiotemporal control. To circumvent these challenges, we used single-cell RNA-sequencing (scRNA-seq) analysis. Since airway smooth muscle differentiates continuously during lung branching morphogenesis^3^, scRNA-seq analysis at a single snapshot in time can be used to interrogate cell identities ranging from undifferentiated progenitors to nascent smooth muscle cells to mature airway smooth muscle^26,27^.

scRNA-seq analysis of *E*11.5 mouse lungs revealed a continuum of mesenchymal and smooth muscle cell identities, but no distinct smooth muscle progenitor population. Instead, mesenchymal cell heterogeneity reflects spatially distinct, Wnt-dependent sub-epithelial and sub-mesothelial compartments. Live-imaging and tension-sensor experiments revealed differences in motility and cortical tension between compartments, and showed that airway smooth muscle cells arise from the sub-epithelial mesenchyme. Computationally reconstructing this differentiation trajectory showed that cytoskeletal, adhesion, and Wnt signaling genes are upregulated early during differentiation, and that the airway smooth muscle gene-expression program is largely *Myocd*-independent. Our computational analyses uncovered a role for Yap1 in mesenchymal patterning and showed that differentiating cells systematically downregulate genes involved in proliferative metabolism. Finally, we found that manipulating Wnt signaling affects the enrichment of nascent smooth muscle cells and F-actin in the mesenchyme, thus altering mesenchymal constraint on the epithelium and influencing epithelial branching. Overall, we provide the first single-cell view of airway smooth muscle differentiation and demonstrate that the earliest steps of this cascade involve Wnt-dependent mesenchymal stiffening that shapes epithelial branches.

## Results

### scRNA-seq analysis reveals expected populations of cells in the embryonic murine lung

We carried out scRNA-seq using cells isolated from the left lobes of lungs of CD1 mouse embryos harvested at *E*11.5 and analysed the data with Seurat^28^ (**Fig. 1a, Supplementary Fig. 1a-j**). After removing contaminating cells and correcting for cell-cycle stage (**Supplementary Fig. 1k-o**), clustering identified 6 distinct populations, each containing cells from both replicates (**Fig. 1b-c**). We identified mesenchymal clusters expressing homeobox B6 (*Hoxb6*) and pleiotrophin (*Ptn*), a smooth muscle cluster expressing *Acta2*, an epithelial cluster expressing thyroid transcription factor 1 (TTF-1 or *Nkx2.1*), a vascular-endothelial cluster expressing VE-cadherin (*Cdh5*), and a mesothelial cluster expressing Wt1 (**Fig. 1b-e, Supplementary Table 1**).

**Figure 1.**
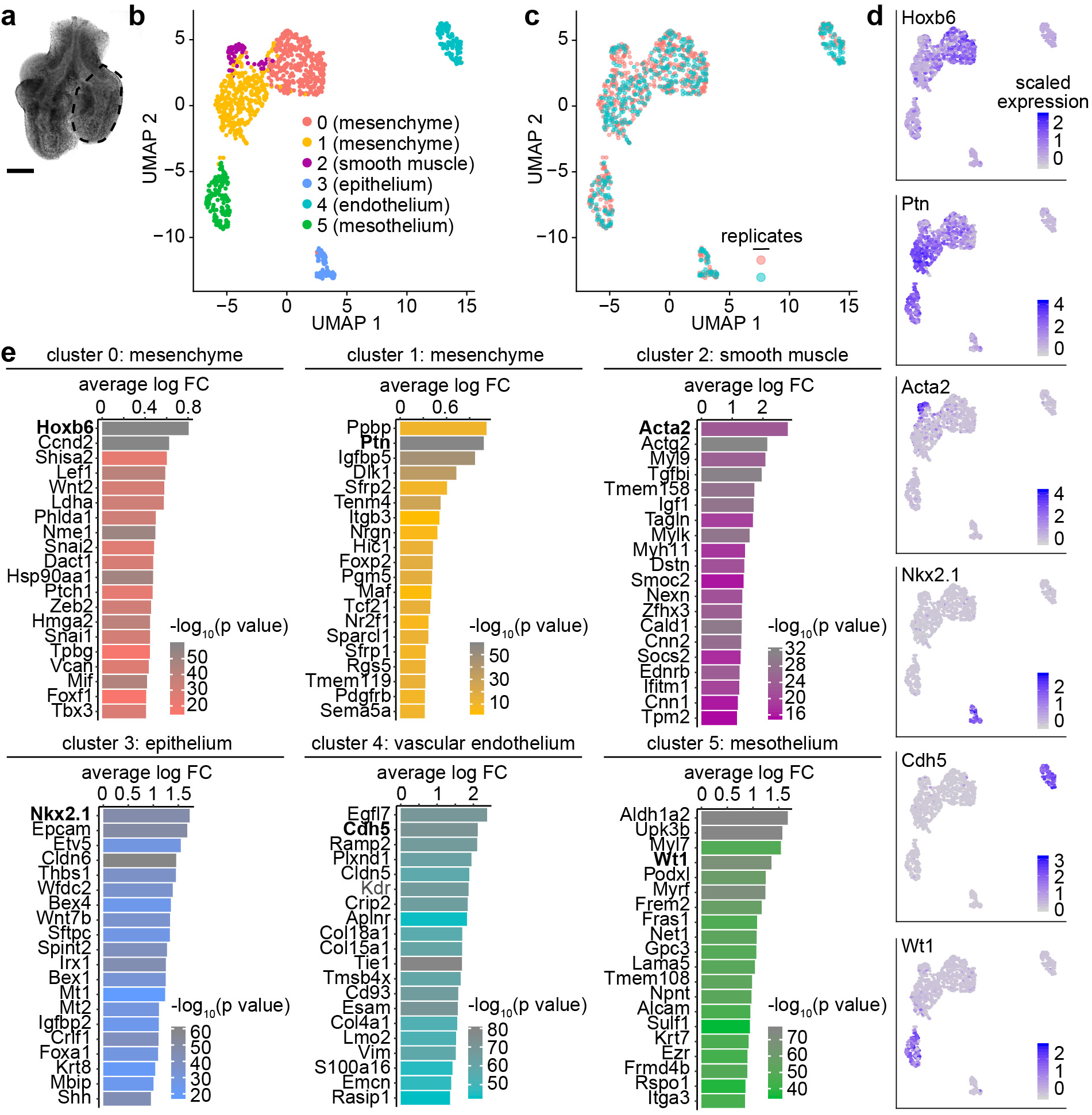
Clustering scRNA-seq data from *E*11.5 mouse lungs reveals expected cell populations. (**a**) Image of sample lung used for scRNA-seq experiment. Each lung was micro-dissected to isolate the left lobe (indicated by dotted outlines). Scale bar shows 100 µm. (**b-c**) UMAP plots of cell cycle-corrected embryonic lung cells isolated by scRNA-seq of *E*11.5 CD1 mouse lungs, color-coded either by cluster (b) or by replicate (c). (**d**) UMAP plots from (b-c) color-coded to show the expression of population-specific genes. (**e**) Top 20 markers for each cluster shown in Fig. 1b based on log fold-change (FC) and color-coded by adjusted p-value.

We then isolated and reclustered the closely-related mesenchymal and smooth muscle populations, and found that the *Hoxb6-*, *Ptn-*, and *Acta2*-expressing cells still clustered together in a similar pattern (**Supplementary Fig. 2a-b**). Previous studies used lineage-tracing of Axin2-, Fgf10-, Gli1-, and Wt1-expressing cells in an attempt to identify smooth muscle progenitors^24,25^. However, none of these markers labeled a distinct population in our dataset (**Supplementary Fig. 2d-g**). Further analysis of reclustered mesenchymal cells provided no additional evidence for a transcriptionally distinct smooth muscle progenitor (**Supplementary Fig. 2h-q**). Similar to other developing tissues^29,30^, the embryonic pulmonary mesenchyme does not harbor a distinct progenitor population, but contains of a continuum of cell states.

### Undifferentiated mesenchymal cell clusters represent spatially distinct populations

To gain insight into mesenchymal patterning, we examined the expression of markers that distinguished the mesenchymal clusters 0 and 1 (**Supplementary Table 2**). We chose markers differentially expressed between the two clusters for which antibodies are available (Lef1 and Foxp1). Immunofluorescence analysis of *E*11.5 lungs showed that Lef1^+^ cells are present adjacent to the epithelium, suggesting that these sub-epithelial cells may represent cluster 0 (**Fig. 2a**). Conversely, Foxp1^+^ cells are mainly in sub-mesothelial regions and in the medial mesenchyme, suggesting that these cells comprise cluster 1 (**Fig. 2a**). Quantification of intensity profiles of Lef1 as a function of distance from the epithelium and of Foxp1 as a function of distance from the mesothelium showed that these markers are present in opposing gradients across the mesenchyme (**Fig. 2b**). We found similar patterns of Pitx2, an alternate marker for cluster 1 (**Supplementary Fig. 3a-c**). Apart from their differential expression patterns in the mesenchyme, Lef1 and Foxp1 are detected in smooth muscle (**Fig 2c, Supplementary Fig. 3d-e**), and Foxp1 is detected in the epithelium and mesothelium (**Fig 2a, c**). These data suggest that the computationally identified mesenchymal clusters represent spatially distinct populations within the embryonic pulmonary mesenchyme. Therefore, we delineated two spatially distinct mesenchymal compartments, hereafter referred to as sub-epithelial (cluster 0) and sub-mesothelial (cluster 1), following previously established nomenclature^16,17^ (**Fig. 2d**). The observed gradients of Lef1^+^ and Foxp1^+^ cells, and the fact that a portion of mesenchymal cells express both markers (**Supplementary Fig. 3f**), suggests that mesenchymal cells exist in a continuum between these two compartments rather than as two separate cell types.

**Figure 2.**
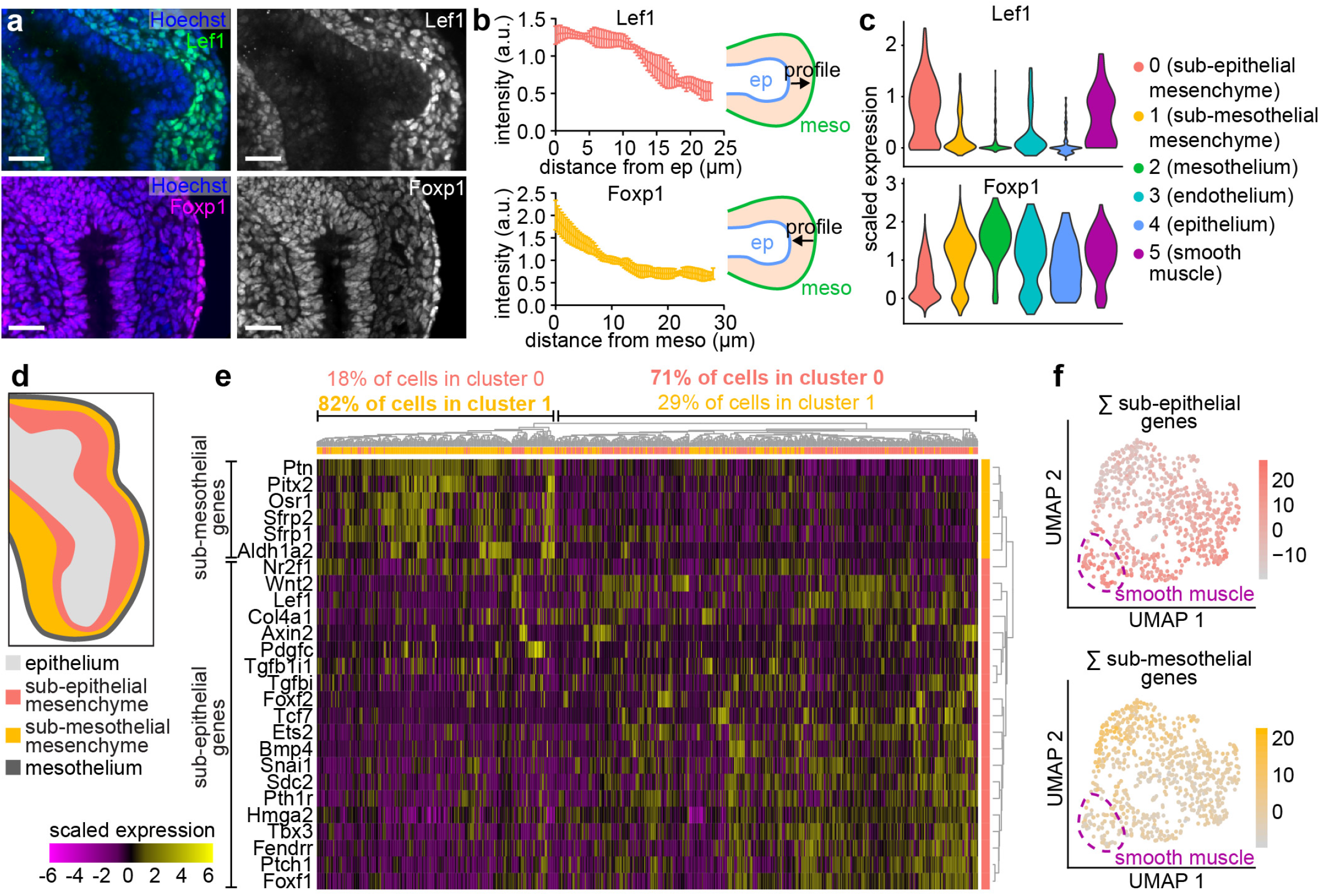
Computationally identified mesenchymal clusters represent spatially distinct populations. (**a**) *E*11.5 lungs immunostained for cluster 0 marker Lef1 or for cluster 1 marker Foxp1 and counterstained with Hoechst. Scale bars show 25 µm. (**b**) Quantifications of Lef1 and Foxp1 intensity profiles emanating from the epithelium (for Lef1) or from the mesothelium (for Foxp1). Schematics show lines and direction along which intensity profiles were measured. Mean and s.d. are plotted (n=4). (**c**) Violin plots showing the expression of *Lef1* or *Foxp1* in each cluster. (**d**) Schematic depicting the sub-epithelial and sub-mesothelial compartments of the mesenchyme. (**e**) Heatmap showing the expression of genes specific to either mesenchymal compartment. Genes (rows) are clustered based on the dendrogram to the right. Cells (columns) are clustered based on the dendrogram above, and each column is color-coded according to original cluster identity from Fig. 1b. (**f**) UMAP of mesenchymal and smooth muscle cells color-coded according to the sum of their expression of either sub-epithelial or sub-mesothelial mesenchyme marker genes. Dotted line indicates location of smooth muscle cells.

To further validate these conclusions, we took advantage of published in situ hybridization results available via the Mouse Genome Informatics (MGI) database. We downloaded a list of genes that are detected in the pulmonary mesenchyme from *E*11.5 to *E*12.5 and classified them by expression pattern (**Supplementary Table 3**). We then focused on those that were clearly sub-epithelial or sub-mesothelial and that were detected at sufficient levels in our scRNA-seq dataset. Unsupervised clustering of mesenchymal cells based on their expression of region-specific genes showed that sub-epithelial cells group together and have similar expression of sub-epithelial markers and, likewise, that sub-mesothelial cells group together and have similar expression of their corresponding markers (**Fig. 2e**). UMAPs color-coded according to the summed expression of either sub-epithelial or sub-mesothelial genes show that the two sets of markers are expressed by cells in complementary regions of the UMAP (**Fig. 2f**). These findings are consistent with our conclusion that the computationally identified clusters represent spatially distinct compartments.

We next asked whether these mesenchymal compartments persist over the pseudoglandular stage of lung development. At *E*12.5 and *E*14.5, we detected Lef1^+^ cells in the sub-epithelial mesenchyme around branch tips and Foxp1^+^ cells in the sub-mesothelial mesenchyme (**Supplementary Fig. 3f-i**). Individual Foxp1^+^ cells are also visible in the mesenchyme in between branches, where they likely represent vascular endothelial cells (**Fig. 2c, Supplementary Fig. 3g, i**). Therefore, the lung maintains a sub-epithelial, Lef1^+^ mesenchymal population and a sub-mesothelial, Foxp1^+^ mesenchymal population over the course of branching morphogenesis. The mesenchyme in between branches (with low Lef1 and Foxp1 immunostaining) may take on new characteristics, perhaps to support formation of vasculature.

### Wnt signaling regulates cell identity in the embryonic pulmonary mesenchyme

We hypothesized that the mesenchymal populations identified in our scRNA-seq dataset represent cell types with important functions in branching morphogenesis. Indeed, GO enrichment analysis of genes upregulated in each mesenchymal cluster revealed that both clusters are enriched for genes involved in “morphogenesis of a branching epithelium” (**Fig. 3a, Supplementary Tables 2, 4**). Additionally, our analysis revealed a significant enrichment of genes with GO terms related to Wnt signaling. We therefore examined differentially-expressed Wnt signaling-related genes (**Fig. 3b**). Among the genes upregulated in the sub-epithelial cluster were *Lef1*, a Wnt signaling effector^31^, and *Wnt2*, a Wnt ligand implicated in airway smooth muscle differentiation^23^. Overall, the sub-epithelial cluster expresses more activators and targets of Wnt signaling, while the sub-mesothelial cluster expresses more inhibitors. Each mesenchymal cluster is therefore characterized by a distinct Wnt-signaling signature and, consistent with previous studies^19,20,32^, the sub-epithelial cluster appears to be more Wnt-active.

**Figure 3.**
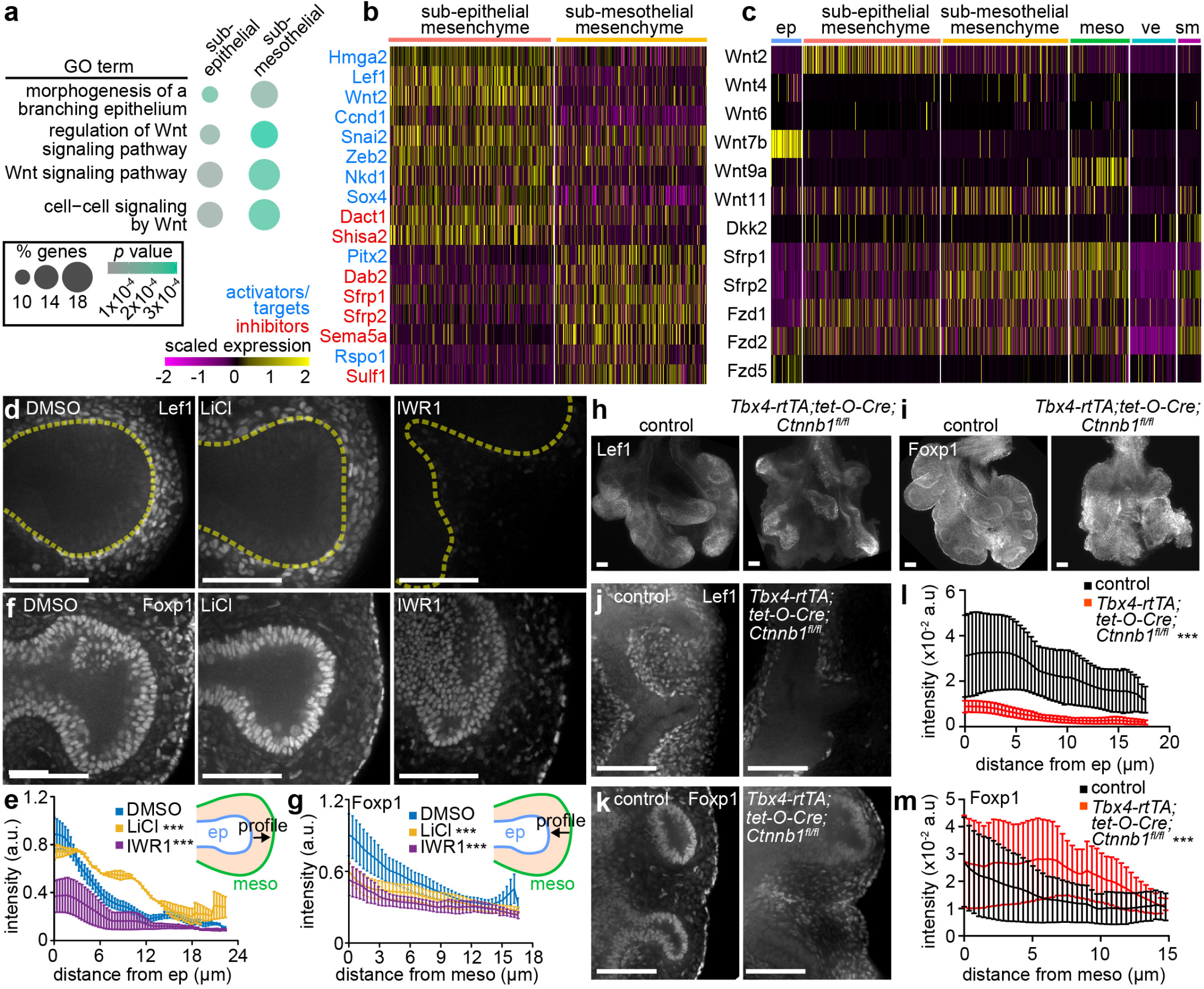
Wnt signaling regulates cell identity in the embryonic pulmonary mesenchyme. (**a**) Bubble plot showing the enrichment percentage and adjusted p-value of relevant GO terms identified based on genes upregulated in each mesenchymal compartment. (**b**) Heatmap showing the expression of Wnt-associated genes upregulated in either mesenchymal cluster. Activators and targets are colored in green, inhibitors are colored in orange. (**c**) Heatmap showing the expression of Wnt ligands, secreted inhibitors, and receptors detected in either mesenchymal cluster and in clusters containing cells from the mesothelium (meso), vascular endothelium (ve), epithelium (ep), and smooth muscle (sm). (**d-g**) Confocal sections and quantifications of Lef1 and Foxp1 intensity profiles around branch L.L2 in lungs isolated at *E*11.5 from CD1 embryos and immunostained for Lef1 or Foxp1 after treatment with either DMSO, LiCl (10 mM), or IWR1 (100 µM) for 24 hr (n=2-6). Yellow dashed lines indicate border of the epithelium. Schematics show lines and direction along which intensity profiles were measured. Mean and s.e.m. are plotted, and curves were compared using two-way ANOVA. (**h-m**) *E*12.5 control and *Tbx4-rtTA;tet-O-Cre;Ctnnb1^fl/fl^* lungs immunostained for Lef1 or Foxp1 and quantifications of Lef1 and Foxp1 intensity profiles (n=3). Low-magnification z-projections (h, i) and high-magnification confocal slices (j, k) are shown. Scale bars show 50 µm. *** indicates *p*<0.0001.

Signals from the epithelium and mesothelium have previously been shown to regulate behaviors of the sub-epithelial and sub-mesothelial mesenchyme, respectively^16,17^. We therefore hypothesized that Wnt signals from these tissues could establish compartments of differential Wnt signaling across the epithelium-to-mesothelium axis. Tissue-specific analysis using our scRNA-seq dataset confirmed that the epithelium expresses *Wnt7b* and the sub-epithelial mesenchyme expresses *Wnt2*^19^ (**Fig. 3c**). The sub-mesothelial mesenchyme expresses *Wnt11*, the mesothelium expresses *Wnt9a*, and both tissues express the inhibitors *Sfrp1* and *Sfrp2*, suggesting that Wnt signaling may be inhibited at the mesothelial end of the axis (**Fig. 3c**). Vascular-endothelial and smooth muscle cells do not show any striking expression patterns of Wnt ligands or secreted inhibitors. *Fzd1* and *Fzd2* are the only Wnt receptors detected in the mesenchyme, but they are not preferentially enriched in either cluster (**Fig. 3c**), suggesting that differential Wnt signaling across the mesenchyme arises from patterned expression of ligands and inhibitors, rather than from differential expression of receptors.

To determine whether Wnt signaling can regulate the compartments of the pulmonary mesenchyme, we manipulated this pathway in lungs explanted from *E*11.5 CD1 mouse embryos. To activate Wnt signaling, we inhibited GSK3 by treating lung explants with LiCl^20^, which maintains nuclear β-catenin levels and increases cytoplasmic Axin2 intensity in the mesenchyme (**Supplementary Fig. 4a-d**). Conversely, to downregulate Wnt signaling, we treated explants with IWR1, which stabilizes Axin2 leading to enhanced activity of the β-catenin destruction complex^20^. As expected given its mechanism of action, IWR1 treatment does not affect Axin2 intensity, but reduces levels of nuclear β-catenin in the mesenchyme (**Supplementary Fig. 4a-d**).

We then quantified the domains of Lef1^+^ sub-epithelial cells and Foxp1^+^ sub-mesothelial cells. We plotted Lef1-intensity profiles as a function of distance from the tip of the epithelial branch L.L2 (the second domain branch of the left lobe, which initiates and extends during the culture period^9,33^) and Foxp1-intensity profiles as a function of distance from the mesothelium. Inhibiting Wnt by treating with LiCl resulted in an expanded domain of Lef1^+^ cells around epithelial branch tips and fewer Foxp1^+^ cells than DMSO-treated controls (**Fig. 3d-g**). Conversely, activating Wnt by treating with IWR1 resulted in a significant reduction in Lef1 intensity and a random distribution of Foxp1^+^ cells in the mesenchyme adjacent to the tip of branch L.L2 (**Fig. 3d-g**). These changes occurred without significant effects on the number of mesenchymal cells between the epithelium and mesothelium (**Supplementary Fig. 4e**). Overall, activating Wnt signaling expands the sub-epithelial mesenchyme at the expense of the sub-mesothelial mesenchyme, and inhibiting Wnt shrinks the sub-epithelial compartment.

To confirm our findings with a genetic approach, we deleted *Ctnnb1* (β-catenin) from the pulmonary mesenchyme using *Tbx4-rtTA;tet-O-Cre*. We found that inhibiting Wnt signaling genetically leads to a dramatic reduction of the Lef1^+^ sub-epithelial mesenchymal compartment (**Fig. 3h, j, l**), in line with previous work^21^, and a corresponding expansion of the Foxp1^+^ sub-mesothelial compartment (**Fig. 3i, k, m**). Consistently, others have shown that enhancing Wnt signaling via mesenchymal deletion of the Wnt-inhibitor *Apc* (Adenomatous polyposis coli) results in increased expression of the sub-epithelial mesenchymal marker *Vcan* (versican; **Supplementary Table 2**)^22^. Our predictions based on scRNA-seq analyses are therefore confirmed by both pharmacological and genetic manipulations.

To determine whether mesenchymal patterning also depends on Shh, we treated explants with the Shh antagonist cyclopamine, which prevents smooth muscle differentiation and decreases sub-epithelial mesenchymal cell proliferation^9,16^. In the presence of cyclopamine, the Lef1^+^, sub-epithelial domain is reduced around branch tips, but the Foxp1^+^ sub-mesothelial domain is slightly expanded (**Supplementary Fig. 4f-i**). We then asked whether Wnt or Shh could override the compartment-specific effects of Shh, and vice versa. We combined cyclopamine and LiCl treatments and found that activating Wnt signaling rescues the loss of Lef1^+^ cells caused by Shh inhibition (**Supplementary Fig. 4j, l**). However, Foxp1 intensity is much lower than controls and similar to LiCl treatment alone (**Supplementary Fig. 4k, m**), suggesting that the positive effects of Shh inhibition on the sub-mesothelial compartment are indirect and cannot overcome the effects of activated Wnt signaling.

### Mesenchymal cell motility and cortical forces are spatially patterned during airway branching morphogenesis

Our observations thus far suggest that the two mesenchymal clusters represent a continuum of cell states that can be shifted by Wnt signaling, instead of two distinct tissues. We therefore investigated whether mesenchymal cells transition between compartments and/or into the airway smooth muscle lineage. We used *Dermo1-Cre-*driven expression of *R26R-Confetti* to fluorescently label mesenchymal cells for time-lapse imaging analysis of lung explants isolated at *E*11.5 (**Fig. 4a**). *Dermo1-Cre* is active throughout the pulmonary mesenchyme^16^, including in Lef1^+^ sub-epithelial and Foxp1^+^ sub-mesothelial mesenchymal cells and the smooth muscle layer (**Supplementary Fig. 5a-d**). During time-lapse imaging, Confetti-labelled cells elongate around branching epithelial buds (**Fig. 4a**). Strikingly, Confetti-labelled cells are not clustered together, as has been observed in other tissues labeled with Confetti^34^, suggesting that embryonic pulmonary mesenchymal cells rearrange extensively as the airway epithelium undergoes branching morphogenesis (**Fig. 4a**). Indeed, mesenchymal cells are suprisingly motile (**Supplementary Video 1**) and exhibit a range of migratory behaviors, from sub-diffusive to directed (**Fig. 4b, Supplementary Fig. 5e-j**), as revealed by cell tracking and mean squared displacement (MSD) analyses. Migration speeds are highly variable and are greatest among sub-epithelial mesenchymal cells (**Fig. 4b-c**), demonstrating that cell motility differs between mesenchymal compartments.

**Figure 4.**
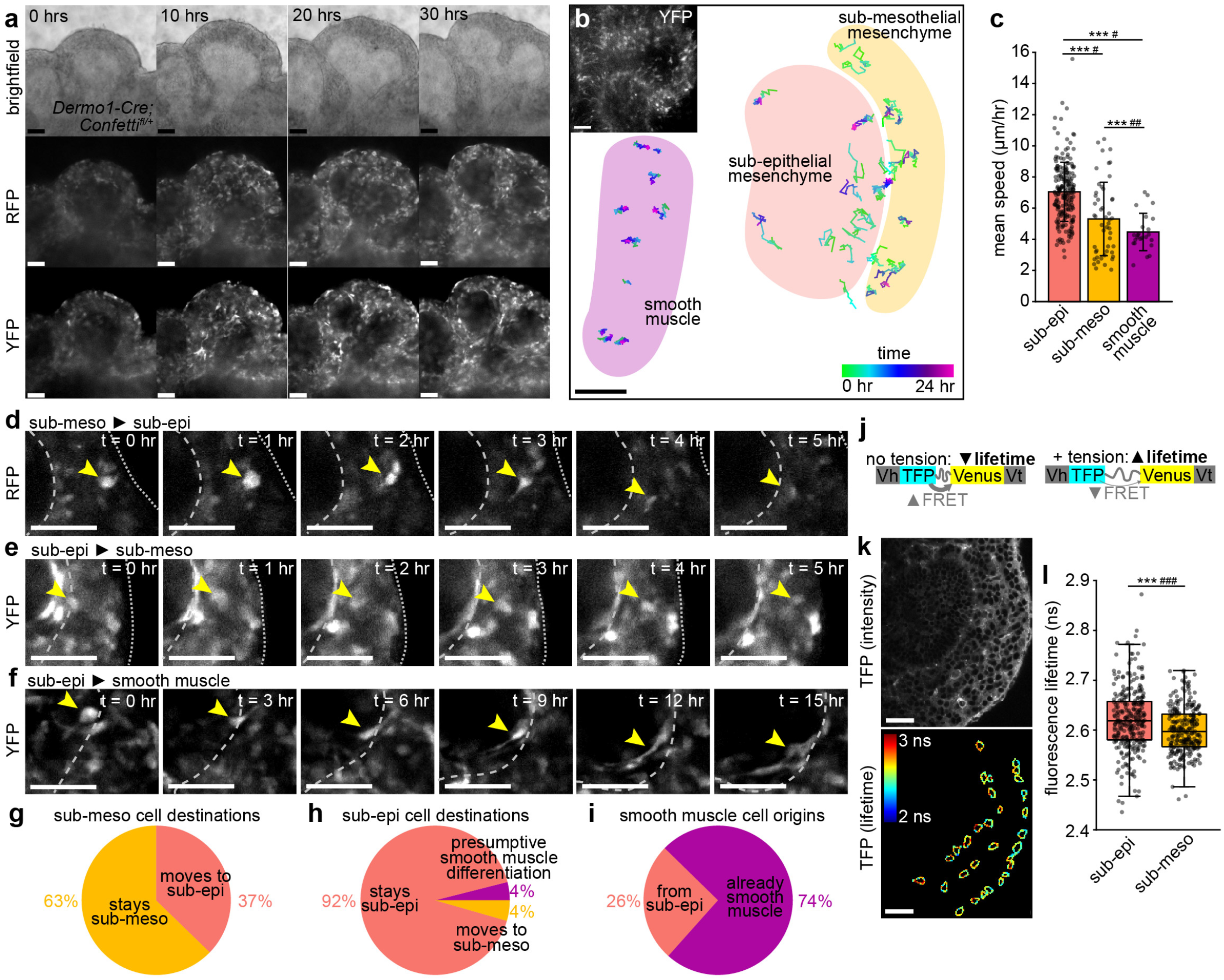
Embryonic pulmonary mesenchymal cells exhibit spatially patterned migratory behaviors and cortical tension. (**a**) Snapshots from time-lapse analysis of *Dermo1-Cre;Confetti^fl/^*lungs isolated at *E*11.5 showing a bifurcating epithelial branch surrounded by sparsely fluorescently-labelled mesenchymal cells. (**b**) Plot of cell trajectories from tracking time-lapse analysis of *Dermo1-Cre;Confetti^fl/^* lungs isolated at *E*11.5. Tracks are color-coded based on time-point. Shaded areas show mesenchymal compartments and smooth muscle regions. Inset in top left corner shows a snapshot of the time-lapse from which the cell tracks are taken. (**c**) Mean speed of tracked cells of each mesenchymal cell type (n=6 time-lapses, n=280 cells total). Error bars show s.d. (**d-f**) Snapshots from time-lapse analyses of *Dermo1-Cre;Confetti^fl/^* lungs isolated at *E*11.5 showing a mesenchymal cell migrating from the sub-mesothelial to the sub-epithelial mesenchyme (d) and vice versa (e), and showing a sub-epithelial mesenchymal cell elongating and presumably differentiating into smooth muscle (f). Yellow arrowhead indicates tracked cell, dashed line indicates the epithelium and dotted line indicates the mesothelium. (**g-h**) Percentage of tracked cells from the sub-mesothelial (g; *n* = 51) or the sub-epithelial (h; *n* = 203) mesenchyme that stay within their original compartment, cross compartments, or elongate, presumably differentiating into smooth muscle. (**i**) Percentage of tracked smooth muscle cells that were already smooth muscle or that were observed to differentiate from sub-epithelial mesenchymal cells (*n* = 26). (**j**) Schematic of VinTS. Vh is vinculin head, Vt is vinculin tail. (**k**) Fluorescence intensity of VinTS in *E*12.5 lung and lifetime of TFP donor along mesenchymal cell edges. (**l**) Fluorescence lifetime of sub-epithelial and sub-mesothelial mesenchymal cells (n=6 samples, n=261-282 cells). Box shows 25^th^ and 75^th^ percentiles, center line shows median. Scale bars show 50 µm. Groups were compared using two-sided t-test and Bartlett’s test for unequal variances. *** indicates *p*<0.0001 for t-test and ^#^ indicates *p*<0.05, ^##^ indicates *p*<0.001, and ^###^ indicates *p*<0.0001 for Bartlett’s test.

We observed that mesenchymal cells can move between compartments over timescales that are short (∼5 hours) relative to that of branching in culture (∼20 hours). The majority of cells remain within their compartments over the timescales analyzed, but a fraction cross over into the opposite compartment (**Fig. 4d-e, g-h**), possibly contributing to the continuum of transcriptional states observed in the mesenchyme. Most smooth muscle cells that we tracked were already elongated around the epithelium, but some originated from rounder, sub-epithelial cells at branch tips (**Fig. 4f, i**). Consistent with previous conclusions from fixed tissues^18^, our time-lapse data show that airway smooth muscle cells arise from the sub-epithelial mesenchyme.

Cell migration and rearrangements are regulated by cortical tension, which can be estimated using the FRET-based vinculin tension sensor (VinTS)^35^ (**Fig. 4j**). A VinTS knock-in mouse was recently generated and validated to infer cortical forces during mandibular arch morphogenesis^36^. Live-imaging and fluorescence lifetime imaging microscopy (FLIM) of *E*12.5 VinTS lungs revealed that the epithelium and sub-epithelial mesenchyme have the longest average fluorescence lifetimes (highest tension), while the vasculature has the lowest (**Supplementary Fig. 5k-l**). Consistent with the observed differences in migratory behavior^36^, we found that sub-epithelial mesenchymal cells have higher and more variable (indicative of population-level heterogeneity and/or fluctuations within individual cells) fluorescence lifetimes than sub-mesothelial mesenchymal cells (**Fig. 4k-l**). These data indicate that sub-epithelial cells experience higher cortical tension. We therefore conclude that the two transcriptionally-distinct mesenchymal compartments have different migratory behaviors and mechanical properties.

### Sub-mesothelial mesenchymal cells may give rise to vascular smooth muscle

Our data show that the sub-epithelial mesenchyme gives rise to airway smooth muscle, but computational clustering suggests that the sub-mesothelial mesenchyme is also closely related to smooth muscle. We therefore hypothesized that the sub-mesothelial mesenchyme could give rise to vascular smooth muscle cells. Mature vascular smooth muscle is not yet present in the lung at *E*11.5^37^, but there could be cells in early stages of differentiation down this lineage. To test the plausibility of this hypothesis, we examined the levels of markers enriched in either airway or vascular smooth muscle^3^ (**Supplementary Fig. 6a**). The expression of airway smooth muscle markers is highest in the smooth muscle cluster (**Supplementary Fig. 6b-f**). Among these, *Foxf1*, *Mylk* and *Nog* are enriched in sub-epithelial compared to sub-mesothelial mesenchyme, as expected given that sub-epithelial mesenchyme gives rise to airway smooth muscle. Conversely, the vascular smooth muscle markers *Heyl* and *Speg* are slightly enriched in the sub-mesothelial mesenchyme (**Supplementary Fig. 6g-h**), suggesting that this compartment might contain vascular smooth muscle precursors. Reclustering of smooth muscle cells alone did not reveal a distinct vascular smooth muscle population (**Supplementary Fig. 6i-p**). We therefore examined lungs at *E*14.5, when vascular smooth muscle is easily detected around blood vessels, and observed that vascular smooth muscle cells are Lef1^-^ and Foxp1^+^ (**Supplementary Fig. 6q-s**), consistent with our hypothesis that Foxp1^+^, sub-mesothelial mesenchyme may differentiate into vascular smooth muscle. If Foxp1^+^ cells indeed contain vascular smooth muscle progenitors, then an expansion of the Foxp1^+^ domain could lead to increased differentiation of vascular smooth muscle. Consistently, we observed that deleting *Ctnnb1* from the pulmonary mesenchyme, which expands the Foxp1^+^ domain (**Fig. 3j-k**), leads to an increase in vascular smooth muscle (**Supplementary Fig. 6t**).

### Diffusion analysis of mesenchymal cells provides insight into differentiation trajectories

Our scRNA-seq dataset is comprised of undifferentiated mesenchymal cells and mature smooth muscle cells, and theoretically should include cells at every stage of differentiation between these two states. Live-imaging revealed that airway smooth muscle cells originate from the sub-epithelial mesenchyme (**Fig. 4**). We therefore focused on transitions from the sub-epithelial mesenchymal cluster to the smooth muscle cluster. To search for differentiation trajectories, we implemented diffusion analysis using Destiny^27^. Undifferentiated sub-epithelial mesenchymal cells are separated from smooth muscle cells along diffusion component (DC) 1 (**Fig. 5a**). The expression of smooth muscle-related genes is positively correlated with DC1 (**Fig. 5b**). In contrast, gene number, a powerful indicator of cell developmental potential^38^, is negatively correlated with DC1 (**Fig. 5c**), suggesting that progression along DC1 represents cells moving away from a stem-cell-like state and differentiating into airway smooth muscle.

**Figure 5.**
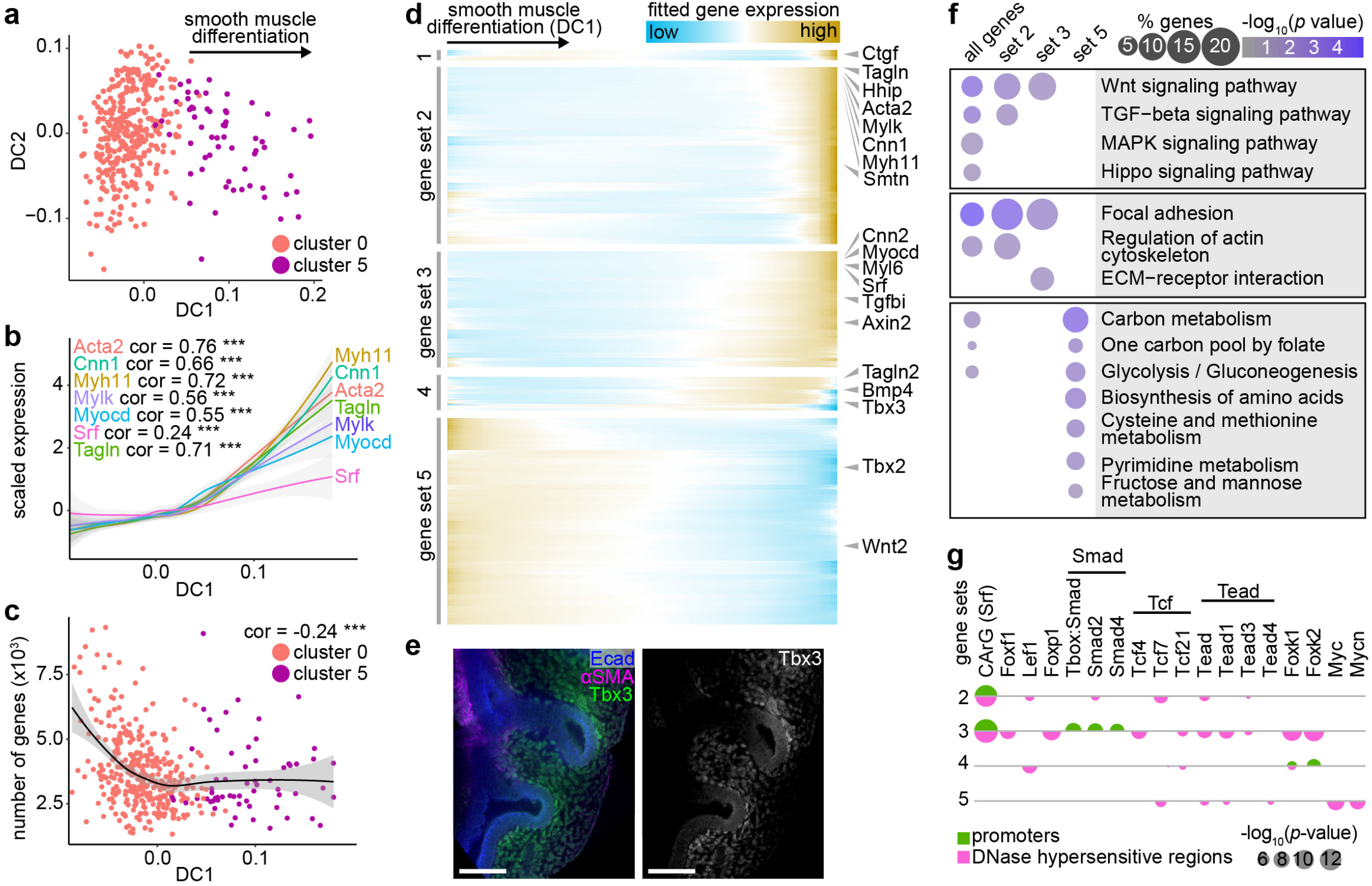
Diffusion analysis reveals differentiation trajectory of mesenchymal cells into airway smooth muscle. (**a**) Diffusion plots of sub-epithelial mesenchymal and smooth muscle cells. (**b**) Scaled expression of known smooth muscle markers versus cell loadings along DC1. (**c**) Number of genes detected per cell versus DC1. (**d**) Heatmap showing 5 gene sets identified using hierarchical agglomerative clustering of spline-fitted gene-expression profiles along DC1. Genes of interest are indicated to the right. (**e**) *E*11.5 lungs immunostained for Tbx3 (gene set 4), Ecad, and αSMA. Scale bars show 50 µm. (**f**) Bubble plot showing the enrichment percentage and adjusted p-value of relevant KEGG pathways for the gene sets defined in e. (**g**) Bubble plot showing the p-values of motifs for transcription factor-binding sites in the promoters of and DNase-hypersensitive regions near genes of sets 2-5. Pearson correlation coefficients and significance are indicated on plots and lines represent smoothed data with standard error shaded in gray. *** indicates *p*<0.0001.

To determine how gene expression evolves over the course of airway smooth muscle differentiation, we fit the expression of each gene along DC1 and applied unbiased clustering to group genes with similar profiles. We focused on four resultant gene sets that increased along DC1 and one that decreased (**Fig. 5d, Supplementary Table 6**). The upregulated gene sets (1-4, numbered from latest to earliest) included genes related to smooth muscle and to Wnt, Shh, TGFβ, Bmp, and Hippo signaling pathways, and the downregulated gene set (5) included the mesenchymal progenitor marker *Tbx2* and the distally-expressed ligand *Wnt2*, suggesting that our analysis had yielded biologically relevant gene sets. *Tbx3* was among the earliest genes activated along the smooth muscle differentiation trajectory, suggesting that it may be a marker for immature smooth muscle cells; consistently, immunofluorescence analysis showed that Tbx3^+^ cells are located in a narrow domain adjacent to the epithelium (**Fig. 5e**), surrounding areas that are not yet wrapped by mature αSMA^+^ smooth muscle.

To identify signaling pathways implicated along the smooth muscle differentiation trajectory, we used KEGG pathway enrichment analysis and carried out motif discovery in promoter and DNase-hypersensitive regions (indicative of accessible chromatin) proximal to genes of each set (**Fig. 5f-g, Supplementary Fig. 7**). Consistent with our hypothesis that DC1 recapitulates smooth muscle differentiation, we found that Srf-binding sites (CArG boxes) are highly enriched in the promoters and DNase-hypersensitive regions of genes in sets 2 and 3 (**Fig. 5g**). Further, we found binding sites for Foxf1, a regulator of airway smooth muscle differentiation^39^, in DNase-hypersensitive regions near genes of set 3 (**Fig. 5g**). We also identified binding sites for the mesenchymal-compartment markers Lef1 and Foxp1 near genes of sets 2, 3, and 4 (**Fig. 5g**).

All gene sets together, and in particular those that increase in expression along DC1, are enriched for genes involved in Wnt, TGFβ, Hippo, Hedgehog, and MAPK signaling (**Fig. 5f**). Accordingly, we found motifs for binding sites of Smad-family transcription factors in promoters of genes in set 3, and of Tcf- and Tead-family members in DNase-hypersensitive regions near genes in sets 2-5 (**Fig. 5g**). Nearly all of these pathways have been implicated in airway smooth muscle differentiation, with the exception of Hippo signaling. To test the role of this signaling pathway, we deleted the Hippo effector *Yap1* from the embryonic mesenchyme using *Dermo1-Cre* and isolated lungs at *E*12.5. Markers for the sub-epithelial and sub-mesothelial compartments are greatly reduced in the mesenchymal knockouts, as is smooth muscle wrapping (**Fig. 6a**), demonstrating that *Yap1* is required for patterning and differentiation in the embryonic pulmonary mesenchyme. Our computational analyses can therefore be used to uncover novel signaling pathways important for lung development.

**Figure 6.**
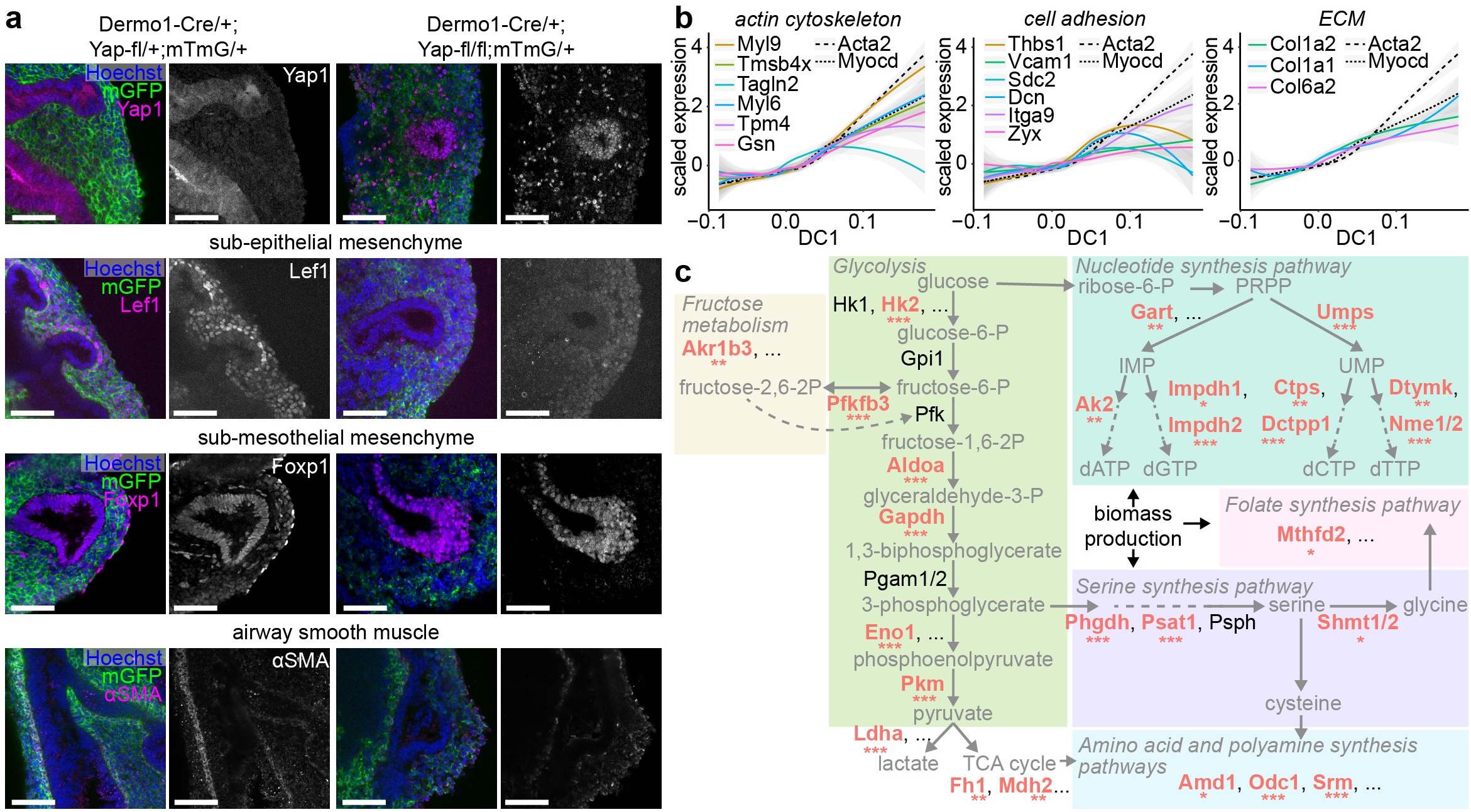
Novel regulators and features of smooth muscle differentiation. (**a**) Sections of *E*12.5 Dermo1-Cre/+;Yap-fl/fl;mTmG/+ lungs and littermate controls immunostained for GFP and either Yap1, Lef1, Foxp1, or αSMA. Yap1^+^ cells in the mesenchyme of mutants are vascular endothelial cells, which are not targeted by *Dermo1-Cre*. (**b**) Scaled expression of genes involved in cytoskeleton, cell adhesion, and extracellular matrix versus cell loadings along DC1 compared to the expression profiles of the smooth muscle markers *Acta2* and *Myocd* (dotted lines). Pearson correlation coefficients and significance are indicated and lines represent smoothed data with standard error shaded in gray. (**c**) Simplified pathway diagram depicting the steps of proliferative metabolism and showing relevant enzymes at each step. Enzymes that are significantly downregulated along DC1 are indicated in bold red font, with significance of spline fit indicated by asterisks. Scale bars show 50 µm. * indicates *p*<0.05, ** indicates *p*<0.001, and *** indicates *p*<0.0001.

Differentiating smooth muscle cells undergo changes in contractility and adhesion to elongate and wrap around the epithelium^3^. Our analyses revealed that genes related to the cytoskeleton, cell adhesion, and the extracellular matrix are upregulated early during differentation, and that their expression accompanies or precedes that of classical smooth muscle markers (*Acta2*) and transcription factors (*Myocd*) (**Fig. 5f, Fig. 6b**). These data suggest that gene expression at the earliest stages of smooth muscle differentiation is *Myocd*-independent. In line with this hypothesis, only a minor fraction (0-6%) of the smooth muscle genes identified using diffusion analysis (**Fig. 5d**) and of smooth muscle cluster markers (11%) (**Fig. 1d**) are differentially expressed in a bulk RNA-seq dataset of *E*13.5 lungs in which *Myocd* is deleted from the mesenchyme^11^ (**Supplementary Fig. 8**). Overall, these data show that the airway smooth muscle gene-expression program is largely *Myocd*-independent.

Finally, we found that genes that decrease in expression along the smooth muscle differentiation trajectory are associated with metabolic pathways (**Fig. 5f**), and that those involved in proliferative metabolism and biomass production are downregulated along DC1 (**Fig. 6c**). Indeed, Myc- and Mycn-binding sites are enriched in DNase-hypersensitive regions near genes in set 5 (**Fig. 5g**), suggesting that proliferative gene expression is downregulated as smooth muscle cells differentiate. Additionally, binding sites for Foxk1 and Foxk2, regulators of glycolysis^40^, are present in the promoters and DNase-hypersensitive regions in gene sets 3 and 4 (**Fig. 5g**). Overall, our analyses uncover a possible role for metabolic reprogramming during smooth muscle differentiation.

### Wnt signaling activates the early stages of smooth muscle differentiation to influence epithelial branching

Our computational analyses suggested the early stages of airway smooth muscle differentation involve changes in cytoskeletal components and Wnt signaling. We therefore evaluated the effects of manipulating Wnt signaling on F-actin accumulation in the mesenchyme around epithelial branch tips and on the expression patterns of early markers of smooth muscle. To visualize smooth muscle cells in the earliest stages of differentiation, we isolated lungs from embryos expressing RFP under the control of the αSMA promoter (SMA-RFP mice). RFP signal can be used to identify nascent smooth muscle cells that do not have sufficient αSMA protein for detection by immunofluorescence analysis. As an alternative marker, we used Tbx3, an early marker of smooth muscle differentiaton identified in our diffusion analyses (**Fig. 5**).

Strikingly, upon activation of Wnt signaling with LiCl treatment, we observed a thick layer of F-actin-rich mesenchyme containing many RFP^+^ cells surrounding the epithelium in LiCl-treated SMA-RFP explants (**Fig. 7a-c**). The domain of Tbx3^+^ nascent smooth muscle cells is also expanded after LiCl treatment (**Supplementary Fig. 9a-b**). Conversely, genetically inhibiting Wnt signaling via mesenchymal deletion of *Ctnnb1* leads to fewer Tbx3^+^ cells around the epithelium and reduced F-actin throughout the mesenchyme (**Fig. 7d-f, Supplementary Fig. 9c-d**).

**Figure 7.**
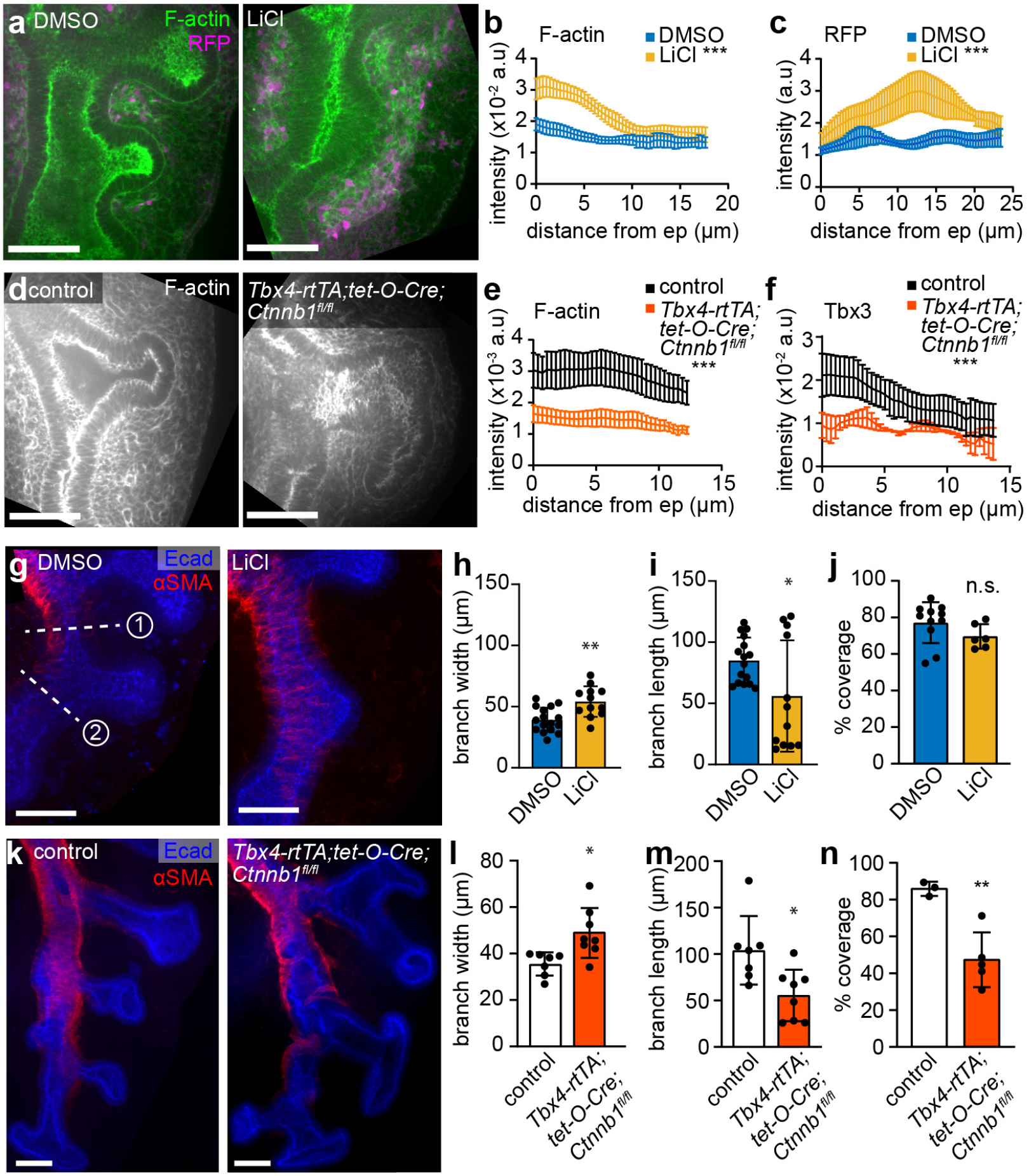
Wnt signaling regulates the accumulation of F-actin and nascent smooth muscle cells in the sub-epithelial mesenchyme to influence epithelial morphology. (**a**) Confocal sections of the left lobe of lungs isolated from *E*11.5 SMA-RFP embryos and immunostained for RFP and labelled with phalloidin (F-actin) after treatment with DMSO or LiCl for 24 hrs. (**b**) Quantification of F-actin intensity profiles emanating from the epithelium (n=3-6). (**c**) Quantification of RFP intensity profiles emanating from the epithelium of lungs isolated from SMA-RFP embryos (n=4-5). (**d**) *E*12.5 control and *Tbx4-rtTA;tet-O-Cre;Ctnnb1^fl/fl^* lungs immunostained for F-actin. (**e**) Quantification of F-actin intensity profiles emanating from the epithelium (n=5). (**f**) Quantification of Tbx3 intensity profiles emanating from the epithelium (n=3). (**g**) Confocal sections of the left lobe of lungs isolated from *E*11.5 CD1 embryos and immunostained for E-cadherin (Ecad) and αSMA after treatment with DMSO or LiCl for 24 hrs. (**h-i**) Width and length of branch L.L2 after 24 hrs of culture (n=13-16). (**j**) Quantification of smooth muscle coverage of the left lobe after 24 hrs of culture (n=6-11). See **Supp. Fig. 9d** for representative segmented images. (**k**) *E*12.5 control and *Tbx4-rtTA;tet-O-Cre;Ctnnb1^fl/fl^* lungs immunostained for Ecad and αSMA. Z-projected images and zoomed-in regions indicated by dashed yellow boxes are shown. (**l-m**) Width and length of branch L.L2 after 24 hrs of culture (n=7-8). (**n**) Quantification of smooth muscle coverage of the left lobe in control and *Tbx4-rtTA;tet-O-Cre;Ctnnb1^fl/fl^* lungs (n=3). See **Supp. Fig. 9e** for representative segmented images. Data in bar graphs were plotted with error bars representing s.d. and were compared using t-test. Intensity profiles were plotted as mean ± s.e.m. and compared using two-way ANOVA. * indicates *p*<0.05, ** *p* < 0.001 and *** *p*<0.0001.

Importantly, these changes to smooth muscle differentiation and mesenchymal F-actin translated to changes in epithelial morphology and smooth muscle wrapping. LiCl treatment impeded branch extension, suggesting that activating Wnt signaling constrains epithelial branching (**Fig. 7g-i**). LiCl treatment does not affect coverage of mature smooth muscle, assessed based on immunofluorescence analysis for αSMA after 24 hours of culture (**Fig. 7g, j, Supp. Fig. 9d**), but the number of nascent smooth muscle cells (that may not yet have robust αSMA expression) is clearly increased (**Fig. 7a, c, Supp. Fig. 9a**). Conversely, genetically inhibiting Wnt signaling in the pulmonary mesenchyme by deleting Ctnnb1 leads to much wider and shorter branches with an irregular shape, suggestive of a lack of constraint (**Fig. 7k-m**). Ctnnb1-mutants also exhibited disorganized αSMA^+^ smooth muscle wrapping and significantly less smooth muscle coverage (**Fig. 7k, l, Supp. Fig. 9e**). These data suggest that Wnt activates the earliest stages of airway smooth muscle differentiation and mesenchymal stiffening, as indicated by local accumulation of F-actin^41,42^, to constrain the branching epithelium (**Fig. 8**).

**Figure 8.**
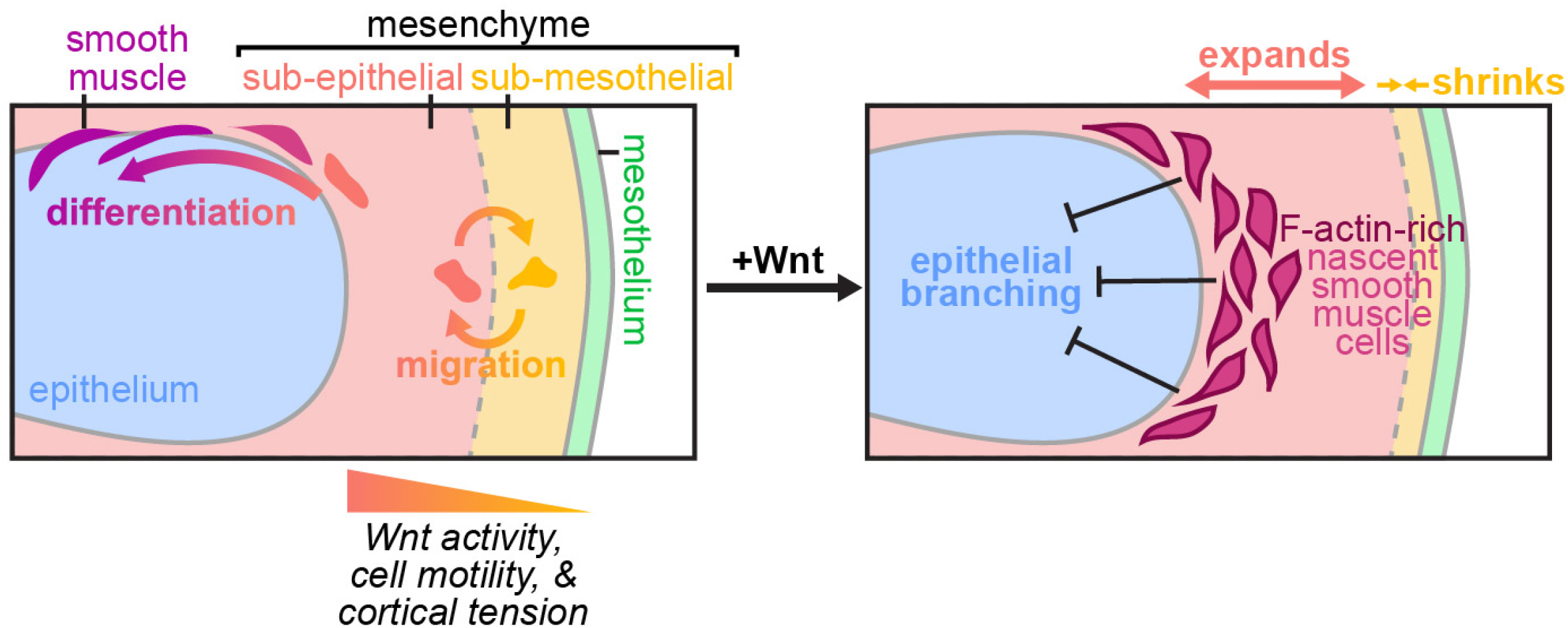
Proposed model for the behaviors and roles of mesenchymal compartments in the embryonic mouse lung. The embryonic pulmonary mesenchyme consists of sub-epithelial and sub-mesothelial compartments that are regulated by Wnt signaling and that are characterized by distinct migratory behaviors and cortical forces. A subset of cells migrate between compartments and sub-epithelial cells give rise to airway smooth muscle. Activating Wnt signaling expands the sub-epithelial compartment, activates the earliest stages of smooth muscle differentiation, and leads to local accumulation of F-actin in the mesenchyme to constrain the branching epithelium.

## Discussion

Here, we provide the first single-cell overview of cell identities and transitions in the embryonic pulmonary mesenchyme during branching morphogenesis of the mouse lung, with a specific focus on airway smooth muscle. scRNA-seq analyses revealed compartment-specific transcriptional heterogeneity across the mesenchyme and uncovered how signaling pathways evolve over the course of airway smooth muscle differentiation. Building off of our computational findings, we identify Wnt signaling as a master regulator of mesenchymal patterning in early lung development and show that Wnt-dependent activation of smooth muscle differentiation and mesenchymal stiffening can physically influence airway epithelial branching, even prior to smooth muscle maturation.

Our data show that the pulmonary mesenchyme contains a continuum of cell identites, which may explain why a distinct marker of smooth muscle progenitors has remained elusive^25^. Unbiased lineage tracing in fixed samples previously showed that new smooth muscle cells at branch tips are members of clones from the adjacent sub-epithelial mesenchyme^18^. Our live-imaging data support these findings, and also reveal a surprising level of cell motility within the embryonic pulmonary mesenchyme. Similar dynamic behaviors have been observed in the developing kidney^41^. The extent and timescales of these rearrangements raise questions about how focal morphogen sources proposed to act as chemoattractants could be maintained as mesenchymal cells migrate and rearrange during epithelial branching morphogenesis^42,43^.

The heterogeneity of mesenchymal cells within developing tissues influences how they respond to morphogenetic signals^44^ and can define if and when they differentiate. Exposing transplanted stalk mesenchyme to a Wnt signal primes it to migrate into a donor tissue and form smooth muscle^18^. It is possible that stalk mesenchyme is comprised of sub-mesothelial mesenchymal cells, and that activation of Wnt signaling converts these cells to a sub-epithelial identity, which we have shown here is associated with greater cytoskeletal tension and motility.

We and others have shown that Fgf9/Shh^16^ and Wnt signaling spanning the epithelium to the mesothelium patterns the embryonic pulmonary mesenchyme. Epithelium-derived Wnt signals also radially pattern smooth muscle differentiation in the ureteric mesenchyme^45^. In the chick intestine, opposing molecular signals from the epithelium and the mesothelium, as well as mechanical forces generated by growth of the intestinal epithelium and contraction of smooth muscle, are integrated to spatiotemporally pattern the differentiation of multiple smooth muscle layers^46^. The role of mechanical forces in regulating pulmonary mesenchymal cell identity have yet to be fully elucidated.

Reconstructing the smooth muscle differentiation trajectory revealed that genes related to the cytoskeleton and cell-matrix adhesion are expressed early during differentiation, prior to and independently of the expression of classical smooth muscle markers. These changes could lead to stiffening of immature smooth muscle cells to regulate diverse morphogenetic events^47–49^. Indeed, we found that Wnt-dependent accumulation of nascent smooth muscle cells and F-actin around emerging branches is associated with local inhibition of epithelial growth. Similarly, activating Wnt signaling in the ureteric mesenchyme leads to ectopic accumulation of smooth muscle progenitors and a hypoplastic ureter^45^. F-actin levels can reflect local tissue stiffness^48,50^, and Wnt signaling can activate actomyosin contractility^51^ and matrix deposition^19^, possibly contributing to tissue stiffening around emerging branches. Our VinTS experiments revealed higher cortical tension in the sub-epithelial mesenchyme, which may allow it to physically sculpt the branching epithelium. Activation of Wnt signaling to promote sub-epithelial mesenchymal fates could enhance this effect.

Our results raise questions about which stage(s) of smooth muscle differentiation and which aspects of smooth muscle cell identity influence branching of the airway epithelium. The smooth muscle gene-expression program extends beyond the influence of a single transcription factor (*Myocd*), suggesting that developing smooth muscle may be robust to the loss or absence of single markers. Indeed, loss of *Acta2* is compensated for by other actin isoforms^12,13^. In the developing lung, *Myocd* deletion decreases the expression of only a few smooth muscle genes (**Supplementary Fig. 8**), and in the developing ureter, expansion of smooth muscle progenitors occurs with minimal increase in the expression of *Myocd*^45^. Consistently, *Myocd* expression is insufficient to drive the entire smooth muscle gene-expression program in cell culture^52^. Combined with previous findings from *Myocd*-knockout animals, our data reveal that the initiation of smooth muscle differentiation is required for airway branching, but mature Myocd^+^ smooth muscle is not.

Our scRNA-seq analyses also uncovered novel regulators of mesenchymal patterning during lung development. We computationally identified and genetically validated a role for Hippo signaling via Yap1 in the embryonic pulmonary mesenchyme. Mesenchymal Yap regulates smooth muscle differentiation around the gastrointestinal epithelium^53^, and crosstalk between Wnt and Yap signaling regulates intestinal stem cell homeostasis^54^. Future investigations will elucidate how Yap influences airway smooth muscle differentiation and whether cooperation with Wnt signaling is involved. Our data also reveal that sub-epithelial cells systematically downregulate enzymes required for proliferative metabolism as they differentiate into smooth muscle, and that Foxk1/2, regulators of glycolysis^40^, may be involved in differentiation.

Metabolic reprogramming is a critical component of stem cell maintenance and differentiation^55^, and enhanced glycolysis acts via Yap to regulate neural crest migration^56^. Notably, Foxk1 interacts with Srf to repress expression of genes including *Acta2*^57^, suggesting that metabolic reprogramming participates in airway smooth muscle differentiation.

In conclusion, our findings reveal the heterogeneity of the embryonic pulmonary mesenchyme and the multitude of signals involved in smooth muscle differentiation. Our computational analyses lay the foundation for future investigations into the mechanisms that regulate cell identity and differentiation in the pulmonary mesenchyme. Further, our approaches could be extended to other organs or organisms to compare tissue patterning and differentiation in diverse developmental contexts.

## Online Methods

### Mice

Breeding of CD1, Ctnnb1^fl/fl^ (JAX 004152), Dermo1-Cre (JAX 008712), R26R-Confetti (JAX 013731), mTmG (JAX 007676), SMA-RFP, Tbx4-rtTA;tet-O-Cre^58^ (gift of Wei Shi), and Yap^fl/fl^ (JAX 027929) mice and isolation of embryos were carried out in accordance with institutional guidelines following the NIH Guide for the Care and Use of Laboratory Animals and approved by Princeton’s Institutional Animal Care and Use Committee. Breeding of VinTS mice^36^ and isolation of embryos were carried out in accordance with the Animals for Research Act of Ontario and the Guidelines of the Canadian Council on Animal Care, and procedures were approved by the Hospital for Sick Children Animal Care Committee. Dermo1-Cre;Confetti^fl/+^ embryos were obtained by mating Confetti homozygous females to Dermo1-Cre heterozygous males. Ctnnb1^fl/fl^ or Yap^fl/fl^ and mTmG mice were bred to generate homozygous Ctnnb1^fl/fl^;mTmG and Yap^fl/fl^;mTmG mice. Dermo1-Cre males were bred to Yap^fl/fl^ females to generate Dermo1-Cre;Yap^fl/+^ males, and these were then mated to Yap^fl/fl^;mTmG females to generate Dermo1-Cre;Yap^fl/fl^;mTmG embryos. Tbx4-rtTA;tet-O-Cre males were bred to Ctnnb1-flox;mTmG females to generate Tbx4-rtTA;tet-O-Cre;Ctnnb1^fl/+^;mTmG males, and these were then mated to Ctnnb1^fl/fl^;mTmG females to generate Tbx4-rtTA;tet-O-Cre;Ctnnb1^fl/fl^;mTmG embryos. Pregnant dams were administered 0.5 mg/ml doxycycline (Sigma) via drinking water from *E*6.5 to *E*12.5, with water replaced every two days. Littermates were used as controls. Pups and embryos were genotyped for Cre, Tbx4-rtTA, mTmG, Ctnnb1^fl^, and/or Yap^fl^ by isolating DNA from the tail snips (pups) or from the head of each embryo, followed by PCR and gel electrophoresis. The forward primer sequence for Cre was GCATTACCGGTCGATGCAACGAGTGATGAG and the reverse primer sequence was GAGTGAACGAACCTGGTCGAAATCAGTGCG. The primer sequences for Tbx4-rtTA were previously published^58^ and the primer sequences for Ctnnb1^fl/fl^, Yap^fl/fl^, and mTmG are provided in the Jackson Labs genotyping protocol for each strain. Where possible, mGFP expression from mTmG was used for genotyping.

### scRNA-seq experiments

Lungs were dissected from CD1 mouse embryos collected at *E*11.5 in cold PBS. Isolated left lobes were grouped by stage, placed in dispase, and mechanically dissociated with tungsten needles. After 10 minutes in dispase at room temperature, DMEM without HEPES and supplemented with 5% fetal bovine serum (FBS, Atlanta Biologicals) was added, and cell suspensions were passed through a filter with 40-µm-diameter pores. The resultant cell suspensions were then processed by the Princeton Genomics Core Facility. The two sample groups were processed separately. Cells were loaded and processed using the Chromium Single Cell 3’ Library and Gel Bead Kit v2 on the Chromium Controller (10X Genomics) following manufacturer protocols. Individual cells were encapsulated in droplets with single gel beads carrying unique barcoded primers and then lysed. Then cDNA fragments were synthesized, barcoded, and amplified by PCR. Illumina sequencing libraries were prepared from the amplified cDNA from each sample group using the Nextra DNA library prep kit (Illumina). The v2 libraries were sequenced on Illumina HiSeq 2500 Rapid flowcells (Illumina) as paired-end 26 + 50 nucleotide reads following manufacturer protocols. Base calling was performed, and raw sequencing reads were filtered using the Illumina sequencer control software to keep only pass-filtered reads for downstream analysis. The 10x CellRanger software version 2.0.1 was used to run the count pipeline with default settings on all FASTQ files from each sample to generate gene-barcode matrices using the *Mus musculus* reference genome mm10-1.2.0.

### scRNA-seq data analysis

scRNA-seq data exported from Cell Ranger were imported into R and processed using the Seurat package^28^. Datasets from all groups were normalized and then integrated based on 2000 variable features (genes) identified by the Seurat function FindVariableFeatures. The integrated dataset was then scaled and analyzed to find neighbors and clusters with the Seurat functions FindNeighbors and FindClusters. Finally, the uniform manifold approximation and projection (UMAP) dimensional reduction was performed to visualize clusters. For initial analyses, all cells were used, which revealed large clusters of blood and immune cells (**Supplementary Fig. 1k-l**). Since these were not the cell populations of interest, we removed them from further analysis (**Supplementary Fig. 1m-n**). We then adjusted the dataset to account for cell-to-cell variability based on cell-cycle stage using the Seurat package. The resulting filtered and adjusted dataset was used for all subsequent analyses. Cluster identities were then assigned based on markers enriched in each cluster that are known to be enriched in each lung cell population. To investigate mesenchymal cells specifically, we repeated the Seurat pipeline outlined above including only cells from clusters 0, 1, and 2 (**Supplementary Fig. 2**). We generated a UMAP, identified markers for each of the 5 new clusters identified, and examined the expression of putative progenitor markers in each cluster (**Supplementary Fig. 2**).

Using published in situ hybridization results available on MGI, we compiled a list of genes that have been detected in the pulmonary mesenchyme at *E*11.5-*E*12.5. Based on MGI entries containing images, we classified the expression pattern of each gene as either sub-epithelial, sub-mesothelial, smooth muscle, everywhere, or weak (**Supplementary Table 3**). Genes that did not fit any of these broad categories because of even more limited expression domains were excluded. A subset of the genes classified as sub-epithelial or sub-mesothelial were detected and included as variable features by the Seurat algorithm. Using the heatmap.2 function in R, we clustered sub-epithelial and sub-mesothelial cells based on their expression of this subset of genes (columns in **Fig. 2e**) and clustered the genes from the curated MGI list (rows in **Fig. 2e**).

To search for functional differences between the mesenchymal clusters identified in **Fig. 1**, we identified genes that were differentially expressed in cluster 0 compared to cluster 1 using the FindMarkers function from the Seurat package, and then grouped these based on whether they were upregulated in cluster 0 or in cluster 1. These gene lists were then subjected to GO enrichment analysis using the clusterProfiler package in R^59^.

### Diffusion analysis of mesenchymal cells

To reconstruct smooth muscle differentiation trajectories, we used the cell-by-gene matrix of 2000 variable genes with scaled expression values generated using the Seurat algorithm and applied the Destiny package in R^27^. Only clusters 0 and 2 were used in this analysis. Three cells from cluster 0 appeared as clear outliers and were excluded from this analysis. To search for groups of genes with similar expression patterns over the course of smooth muscle differentiation, we first used the edgeR package in R to fit the expression of all genes along DC1 using a spline model with three degrees of freedom^60^. The resulting fits were then filtered by adjusted *p* value < 0.05 obtained by implementing a generalized likelihood ratio test with the lrt function from the edgeR package^60^. These fits were then clustered using hierarchical agglomerative clustering with average linkage based on Pearson correlation distance to identify gene sets with similar expression profiles along DC1. Gene sets of interest were then subjected to KEGG pathway enrichment analysis implemented using clusterProfiler^59^.

### Motif discovery

Motif discovery was carried out by examining promoter regions and DNase-hypersensitive regions near genes of interest using a custom Python pipeline and HOMER^61^. Genomic regions based on the mouse ENCODE *E*14 lung DNase-hypersensitivity dataset^62^ were matched to genes from the gene sets identified in **Fig. 5d** based on proximity to their transcriptional start sites. The sequences of these DNase-hypersensitive regions were then extracted and analyzed using the HOMER findMotifs.pl FASTA motif analysis tool^61^ to identify overrepresented motifs compared to a background file of DNase-hypersensitive regions located near the transcriptional start sites of a list of genes randomly generated with the Regulatory Sequence Analysis Tools (RSAT) web interface^63^.

### Organ explant culture and live imaging

Lungs from *E*11.5 CD1 mice were dissected in cold, sterile PBS supplemented with antibiotics (50 units/ml of penicillin and streptomycin) and then cultured on porous membranes (nucleopore polycarbonate track-etch membrane, 8 µm pore size, 25 mm diameter; Whatman) floating on top of DMEM/F12 medium (without HEPES) supplemented with 5% FBS and antibiotics (50 units/ml of penicillin and streptomycin) for 24 hr. Reagents used to manipulate Wnt signaling included LiCl (10 mM; Sigma) and IWR1 (100 μM; Sigma). To inhibit Shh signaling we used cyclopamine (1 μM; Tocris). For live-imaging analysis, Dermo1-Cre/+;Confetti/+ lungs were cultured on Transwell filters (polyethylene terephthalate membrane, 3 µm pore size, 10.5 mm diameter; Corning) within a stage-top incubator (Pathology Devices). Frames were acquired every 30 or 60 min for up to 48 hr under brightfield (1-2 ms exposure per plane for a total of seven planes per time point) or spinning disk confocal illumination (X-light, 122 ms exposure per plane for 5-7 planes per time point) on an inverted microscope (Nikon Ti).

### Vinculin tension sensor experiments

Lungs were isolated from *E*12.5 VinTS embryos and embedded in 1% low melting point agarose on a coverslip fitted into a custom imaging chamber. After an hour of incubation at 37°C, cell culture medium supplemented with 5% FBS was added to the chamber and samples were imaged by confocal and fluorescence lifetime microscopy (FLIM). Imaging and analyses were performed as previously described^36^. Briefly, FLIM was carried out on a Nikon A1R Si laser-scanning confocal microscope with a PicoHarp 300 TCSPC module and a 440 nm pulsed-diode laser (Picoquant). Imaging was performed with a 40×/1.25 *NA* water immersion objective. For each lung, 2 to 3 z-stacks at different locations along the distal left lobe were acquired. Fluorescence lifetime of the donor fluorophore (mTFP1) was estimated using FLIM Fit 5.1.1^64^. Cells of interest were segmented manually in FLIM Fit to obtain an average lifetime per cell. Sub-epithelial mesenchymal cells were at least 2-cell bodies away the from the mesothelium, and sub-mesothelial mesenchymal cells were within 2-cell bodies of the mesothelium.

### Tissue sectioning, immunofluorescence analysis, and imaging

Isolated lungs were fixed in 4% paraformaldehyde in PBS for 15 minutes at room temperature. For sectioning, lungs were washed first in 20% sucrose in PBS, then 30% sucrose in PBS, then left overnight in a 1:1 mixture of OCT and 30% sucrose prior to embedding and freezing in OCT. A Leica CM3050S cryostat was then used to create 10-μm-thick sections for staining on slides. For whole-mount staining and for sections on slides, samples were washed with 0.1% Triton X-100 in PBS and then blocked with 5% goat serum and 0.1% BSA. Samples were then incubated with primary antibodies against αSMA (Sigma a5228, 1:400), Axin2 (Abcam ab32197, 1:200), β-catenin (Sigma SAB4500541, 1:200), E-cadherin (Cell Signaling 3195, 1:200 or Invitrogen 13-1900, 1:200), Foxp1 (Cell Signaling 2005, 1:200), GFP (Invitrogen A-11122, 1:500), Lef1 (Cell Signaling 2230, 1:200), PECAM (Abcam ab28364, 1:200), Pitx2 (gift of Jacques Drouin, 1:300), RFP (Abcam ab62341, 1:400), or Tbx3 (Invitrogen 42-4800, 1:200) followed by incubation with Alexa Fluor-conjugated secondary antibodies (1:200), Alexa Fluor-conjugated phalloidin (1:500) and/or Hoechst (1:1000). Sections on slides were then mounted in Fluorosave. Whole lungs were dehydrated in a methanol series (or isopropanol series for phalloidin-stained lungs) and cleared with Murray’s clear (1:2 ratio of benzyl alcohol to benzyl benzoate). To preserve endogenous fluorescence of Confetti and VinTS lungs, samples were not dehydrated or cleared, but instead mounted in Fluorosave. Confocal stacks were collected using a spinning disk confocal (BioVision X-Light V2) on an inverted microscope (Nikon Ti) using either 20× air, 40× oil, or 60× oil objectives (Nikon).

### Image analysis and statistics

We developed a simple pipeline in MatLab to quantify fluorescence intensity profiles. First, images of Lef1 or Foxp1 immunostaining were background subtracted and then lines were traced either starting at the edge of branch L.L2 and moving outward into the mesenchyme for Lef1 or starting at the mesothelium near branch L.L2 and moving inward into the mesenchyme for Foxp1. Average fluorescence intensity in a 16 × 16 pixel window (5.7 × 5.7 µm) was measured at each point along the line, and the calculation included only the intensity of pixels within cell nuclei based on a mask generated from Hoechst staining. Intensity profiles were averaged from 5 distinct lines per sample, and the resultant curves were compared using two-way ANOVA in GraphPad Prism 5. A similar pipeline was used to quantify Axin2, β-catenin, and F-actin intensity, with small modifications. Axin2 intensity was only averaged from regions with no Hoechst signal, and β-catenin levels were compared as the ratio of mean nuclear (Hoechst^+^) to mean cytoplasmic (Hoechst^-^) pixels. F-actin intensity was measured from all pixels (no mask).

To measure smooth muscle coverage, we first generated masks based on z-projected confocal images of αSMA immunofluorescence. Images were background subtracted and smoothed prior to thresholding. Next, the mask was subject to a “closing” morphological operation in which a binary image is dilated (white regions expand) and then eroded (white regions shrink) in order to close any holes in the initial mask. % coverage was then calculated as the area of the original mask divided by the area of the closed mask (**Supp. Fig. 9d-e**).

### Time-lapse analysis

Cells from time-lapse movies of Dermo1-Cre/+;Confetti/+ embryos were tracked manually in Fiji for as many consecutive time points as possible, and the resultant tracks were analyzed in MatLab. To estimate cell speeds, we first corrected cell tracks for local tissue movement and drift. Since the direction and magnitude of lung growth depends on location within the organ, we corrected cell tracks based on local displacements: for each cell, we computed the mean displacement of all cells within a 150 pixel (111 µm) radius as an estimate of local tissue movement, and then subtracted the cumulative sum of these displacements from the original cell’s track at each time point. This approach allowed us to independently correct for proximal-distal movements near the elongating primary bronchus and lateral movements near expanding branch tips (see tracks for smooth muscle cells vs. sub-epithelial and sub-mesothelial mesenchymal cells in **Supplementary Fig. 5d**). Corrected tracks had slightly smaller instantaneous speeds and much smaller overall displacements (**Supplementary Fig. 5e-f**), indicating that our approach had removed the effects of drift and tissue growth. MSD curves for each tracked cell were generated based on time intervals of up to 7 hours and fit with a 1^st^ degree polynomial to estimate persistence. Only fits with R^2^ greater than 0.4 were included in the comparison between samples. To generate the pie charts in **Fig. 4**, cells were assigned to a tissue compartment (sub-epithelial mesenchyme, sub-mesothelial mesenchyme, or airway smooth muscle) based on their location at the start of their track, and classified as staying within their compartment or crossing into the adjacent compartment by comparing the entire track to tissue growth in the brightfield channel. The boundary between sub-epithelial and sub-mesothelial mesenchyme was between 2- and 3-cell bodies away from the mesothelium. Cells were classified as differentiating into airway smooth muscle based on their positions and their progressively elongating morphology.

### Analysis of bulk RNA-seq of *Myocd*-mutant lungs

Bulk RNA-seq data for 5 control and 5 mutant *E*13.5 lungs in which Myocd had been deleted from the mesenchyme were downloaded from GEO (accession number GSE143394^11^) and processed using the DESeq2 package^65^. Briefly, the DESeq2 package was used to import data, estimate size factors and dispersions, and then calculate differential expression of genes and significance using the Wald test. Log fold changes were shrunk according to the DESeq2 package guidelines. We then used volcano plots to visualize adjusted *p*-values and log_2_ fold changes for either genes that correlated positively (sets 1-4) or negatively (set 5) with DC1 in **Fig. 5d** (**Supplementary Fig. 8a**) or smooth muscle cluster marker genes from **Fig. 1b, d** (**Supplementary Fig. 8c**). Since most of the genes were not significantly differentially expressed, we also plotted adjusted *p*-values against the mean of regularized log transformed counts obtained using the DESeq2 package to confirm that these genes were detected at high enough levels in the bulk RNA-seq dataset and found that there was no bias of differential expression based on transcript abundance (**Supplementary Fig. 8b, d**).

## Abbreviations

DC: diffusion component
Fgf: fibroblast growth factor;
Foxp1: forkhead box P1
Gli1: glioma-associated oncogene 1
Hoxb6: homeobox B6
Lef1: lymphoid enhancer binding factor 1
MGI: Mouse Genome Informatics
Myocd: myocardin
Shh: sonic hedgehog, SMA, smooth muscle actin
SRF: serum response factor
TGF: transforming growth factor
TTF-1: thyroid transcription factor 1
UMAP: uniform manifold approximation and projection
Wnt: wingless-type MMTV integration site family
WT1: Wilms tumor 1

## Acknowledgements

We would like to acknowledge Dr. Wei Wang and the Genomics Core Facility of Princeton University. We thank Dr. Wei Shi (Keck School of Medecine of USC) for generously providing us with the *Tbx4-rtTA;tet-O-Cre* mouse line. We also thank members of the Tissue Morphodynamics Group for helpful discussions and feedback on the mnauscript. This work was supported by an NIH/NICHD R01 (HD0990300), an HHMI Faculty Scholars Award, and an NIH/NHLBI R01 (HL120142) to C.M.N and a CIHR award (MOP 126115) to S.H. K.G. was supported in part by a postgraduate scholarship-doctoral (PGS-D) from the Natural Sciences and Engineering Research Council of Canada and the Dr. Margaret McWilliams Predoctoral Fellowship from the Canadian Federation of University Women. J.M.J. was supported in part by an NIH NRSA Fellowship (F30 HL139039).

## Author Contributions

K.G. and C.M.N. conceptualized the study, designed the experiments, interpreted the data, and wrote the manuscript. K.G. performed the experiments and collected the data. J.M.J. carried out the motif discovery analysis. H.T., M.Z., and S.H. provided all reagents and technical assistance for obtaining VinTS results. All authors provided input on the final manuscript.

