## Supplement for "Patterning the embryonic pulmonary mesenchyme"

**Supplementary Information**

**Supplementary Figure Legends**

**Supplementary Figure 1 – related to Figure 1. scRNA-seq experimental design, summary statistics, and initial clustering.** (**a**) Images of all *E*11.5 lungs used for the two replicates of the scRNA-seq experiment. Each lung was micro-dissected to isolate the left lobe (indicated by dotted outlines). Scale bars show 100 μm. (**b**) Table of general statistics for each sample. UMI (unique molecular identifiers) reflects the number of transcripts captured per cell. (**c-d**) UMI counts as a function of number of barcodes for replicates 1 (c) and 2 (d). Portion of curve above threshold includes all cells (green) and portion below is considered background (grey). (**e**) Table of sequencing statistics for each replicate. Q30 statistics give the percentage of bases in the indicated sequence type with a quality score greater than 30. (**f-g**) Sequencing depth as a function of mean reads per cell for each replicate. (**h-i**) Median genes per cell as a function of mean reads per cell. (**j**) Table of mapping statistics for each replicate. Mapping was performed using STAR. (**k-l**) UMAP plot of all cells isolated by scRNA-seq of *E*11.5 CD1 mouse lungs, color-coded by cluster (k) or by replicate (l). (**m-n**) UMAP plot after all blood and immune cells have been removed, color-coded by cluster (m) or by replicate (n). (**o**) Numbers of clusters by cell type and number of cells in each cluster before and after cell cycle regression.

**Supplementary Figure 2 – related to Figure 1. Pulmonary mesenchymal cells are heterogeneous but ultimately closely related.** (**a-c**) UMAP plots of isolated mesenchymal cells color-coded to show the expression of key markers from Fig. 1: *Hoxb6*, *Ptn*, and *Acta2*. (**d-g**) UMAP from (a) color-coded to show the expression of putative smooth muscle progenitor markers *Axin2*, *Fgf10*, *Gli1*, and *Wt1*. (**h**) UMAP of mesenchymal and smooth muscle cells corrected for cell-cycle stage and color-coded based on cluster identified by Seurat. (**i-m**) Top 20 marker genes for each cluster based on average log fold-change (FC) and color-coded by adjusted p-value. (**n-q**) Violin plots showing the expression of putative smooth muscle progenitor markers *Axin2*, *Fgf10*, *Gli1*, and *Wt1* in each of the mesenchymal clusters represented in (h).

**Supplementary Figure 3 – related to Figure 2. Expression patterns of mesenchymal markers throughout the lung and across developmental stages.** (**a**) *E*11.5 lungs immunostained for cluster 1 marker Pitx2 and counterstained with Hoechst. (**b**) Quantification of Pitx2 intensity profiles emanating from the mesenchyme. Schematic shows lines and direction along which intensity profiles were measured. Mean and s.d. are plotted (n=7). (**c**) Violin plots showing the expression of *Pitx2* in each cluster. (**d-e**) Zoomed-in view of a confocal slice of the left primary bronchus of lungs isolated at *E*12.5 and immunostained for αSMA and either Lef1 or Foxp1. Epithelium indicated by ep. (**f**) Expression levels of Foxp1 plotted against those of Lef1 in mesenchymal cell clusters 0 and 1. (**g-j**) Confocal slices of the left lobe of lungs isolated at *E*12.5 or *E*14.5 and immunostained for either Lef1 or Foxp1 and counterstained with Hoechst. Scale bars show 50 μm.

**Supplementary Figure 4 – related to Figure 3. Manipulating Wnt and Shh signaling ex vivo.** (**a-b**) Sections of lungs isolated at *E*11.5 from CD1 embryos and immunostained for Axin2 or β-catenin and counterstained with Hoechst after treatment with either DMSO, LiCl (10 mM), or IWR1 (100 μM) for 24 hrs. Yellow dashed boxes indicate zoomed-in regions displayed to the right. (**c**) Quantification of mean cytoplasmic Axin2 immunofluorenscence intensity in the mesenchyme for each treatment (n=4-11 sections, n=2-3 lungs). (**d**) Quantification of the ratio of nuclear to cytoplasmic β-catenin immunofluorenscence intensity in the mesenchyme for each treatment (n=3-7 sections, n=2-3 lungs). (**e**) Quantification of the number of mesenchymal nuclei between the epithelium and mesothelium near branch L.L2 for each treatment (n=7-13 lungs). (**f-i**) Confocal sections and quantifications of Lef1 and Foxp1 intensity profiles around branch L.L2 in lungs isolated at *E*11.5 and treated with DMSO or cyclopamine (1 μM) for 24 hr (n=4-5). (**j-m**) Confocal sections and quantifications of Lef1 and Foxp1 intensity profiles around branch L.L2 in lungs isolated at *E*11.5 and treated with DMSO or a combination of cyclopamine and LiCl for 24 hr (n=2-3). Scale bars show 25 μm. Treatment conditions were compared using one-way ANOVA. * indicates $p<0.05$ and *** indicates $p<0.001$

**Supplementary Figure 5 – related to Figure 4. Confetti fluorescent reporters driven by Dermo1-Cre can be used to track single-cell movements in the embryonic pulmonary mesenchyme.** (**a**) Confocal slice of the left lobe of lungs isolated at *E*11.5 from a *Dermo1-Cre;mTmG* embryo and immunostained for E-cadherin and GFP. GFP expression throughout the mesenchyme indicates that Cre is active throughout this tissue. (**b-d**) Confocal sections of *Dermo1-Cre;Confetti*$^{fl/+}$lungs isolated at *E*11.5, cultured for 48 hrs under time-lapse conditions, and immunostained for Lef1, Foxp1, or αSMA and GFP. (**e-f**) Snapshots from a time-lapse of *Dermo1-Cre;Confetti*$^{fl/}$ lung isolated at *E*11.5 (e) and plot of cell tracks (f). Tracks colored in gray are uncorrected and tracks colored in blue are corrected for local tissue movement. Cell-type assignments are indicated by the shaded regions. (**g-h**) Distributions of instantaneous speeds from each cell track (e) and of total cell displacements before and after correcting for local tissue movement. (**i**) Log-log plot of mean MSD curves of each mesenchymal cell type. Error bars show s.d. (**j**) Persistence of cell tracks measured as slope of MSD curves from (g). (**k**) Fluorescence lifetime of each lung tissue. Data points represent average per sample. Box shows $25^{th}$ and $75^{th}$ percentiles, center line shows median. (**l**) VinTS lung stained for PECAM. Cells with high VinTS expression dispersed throughout the mesenchyme are vascular endothelial cells. Scale bars show 50 µm. Groups were compared using two-sided t-test and Bartlett's test for unequal variances. *** indicates $p<0.0001$ and * indicates $p<0.05$ for t-test. $^{\#\#\#}$ indicates $p<0.0001$ and $^{\#}$ indicates $p<0.05$ for Bartlett's test.

**Supplementary Figure 6 – related to Figure 5. Sub-mesothelial cells may preferentially give rise to vascular smooth muscle cells.** (**a**) Table showing the expected expression of each of marker in airway/visceral smooth muscle versus vascular smooth muscle[3]. (**b-h**) Violin plots showing the scaled expression of *Actg2*, *Foxf1*, *Gata5*, *Heyl*, *Mylk*, *Nog*, and *Speg* in each of the mesenchymal and smooth-muscle clusters from **Fig. 1b**. Gene expression per cell cluster compared by Wilcoxon Rank Sum Test using the FindMarkers function. (**i-p**) UMAP of smooth muscle cells computationally isolated from the dataset in **Fig. 1b** color-coded based on scaled expression of marker genes for airway and vascular smooth muscle. (**q-s**) Sections of *E*14.5 lungs immunostained for E-cadherin, αSMA, and either PECAM, Lef1, or Foxp1 and counterstained with Hoechst. Airway smooth muscle (ASM) and vascular smooth muscle (VSM) are indicated. (**t**) *E*12.5 control and *Tbx4-rtTA;tet-O-Cre;Ctnnb1*$^{fl/fl}$ lungs immunostained for αSMA and PECAM. Airway smooth muscle (ASM) and vascular smooth muscle (VSM) are indicated. Scale bars show 50 µm. * indicates $p<0.05$, ** indicates $p<0.001$, and *** indicates $p<0.0001$.

**Supplementary Figure 7 – related to Figure 5. Transcription factors identified in motif discovery analysis are expressed in the embryonic lung.** Motif logos and tissue-specific expression of corresponding transcription factors for results presented in **Fig. 5f**. For expression of Lef1 and Foxp1, see **Fig. 2c**.

**Supplementary Figure 8 – related to Figure 6. The majority of smooth muscle marker genes are Myocd-independent.** (**a**) Volcano plot showing the adjusted p-values and log fold changes of gene-set members (**Fig. 5d**) that were detected in bulk RNA-seq of *E*13.5 Myocd-KO lungs compared to controls. Insets to show each gene set separately with the percentage of genes in each quadrant of the volcano plot indicated. (**b**) Adjusted p-values plotted against the mean of regularized log-transformed counts across all samples showing that genes that were not significantly different were detected at similar levels as genes that were significantly different. Points are color-coded according to gene set and insets to the right show each gene set separately. (**c**) Volcano plot showing the adjusted p-values and log fold changes of smooth-muscle cluster markers (**Fig. 1b, e**) that were detected in bulk RNA-seq of *E*13.5 Myocd-KO lungs compared to controls. The percentages of genes in each quadrant of the volcano plot are indicated. (**d**) Adjusted p-values plotted against the mean of regularized log-transformed counts across all samples.

**Supplementary Figure 9 – related to Figure 7.** (**a-b**) Confocal sections of the left lobe of lungs isolated from *E*11.5 embryos and immunostained for Tbx3 after treatment with DMSO or LiCl for 24 hrs and quantification of Tbx3 intensity profiles emanating from the epithelium (n=3). (**c**) *E*12.5 control and *Tbx4-rtTA;tet-O-Cre;Ctnnb1*$^{fl/fl}$ lungs immunostained for Ecad and Tbx3. (**d-e**) Original and segmented images of lungs immunostained for αSMA used to quantify % coverage of smooth muscle. First, images are processed and thresholded to generate a mask of regions with αSMA staining. Morphological closing (a dilation followed by an erosion) is then performed on the resultant binary images to generate a closed mask that represents the total area of the airways in contact with smooth muscle. % coverage is calculated as the area of the mask ($A_{mask}$) divided by the area of the closed mask ($A_{closed\ mask}$) and reflects the extent to which the airways are wrapped by smooth muscle.

#### Supplementary Tables

**Supplementary Table 1.** Markers for clusters of embryonic lung cells identified in Fig. 1.

**Supplementary Table 2.** Genes that are differentially expressed in either sub-epithelial or sub-mesothelial mesenchymal cells relative to each other.

**Supplementary Table 3.** Curated list of genes from the MGI database that are expressed in the embryonic pulmonary mesenchyme.

**Supplementary Table 4.** GO enrichment analysis results for genes that are differentially expressed in either sub-epithelial or sub-mesothelial mesenchymal cells relative to each other.

**Supplementary Table 5.** Gene sets identified in the diffusion analysis of smooth muscle differentiation in Fig. 5.

**Supplementary Table 6.** KEGG pathway enrichment analysis results for gene sets identified in the diffusion analysis of smooth muscle differentiation in Fig. 5.

#### Supplementary Videos

**Supplementary Video 1.** Time-lapse imaging of lungs isolated from a Dermo1-Cre/+;Confetti/+ embryo at *E*11.5.

### Supplementary Figure 1

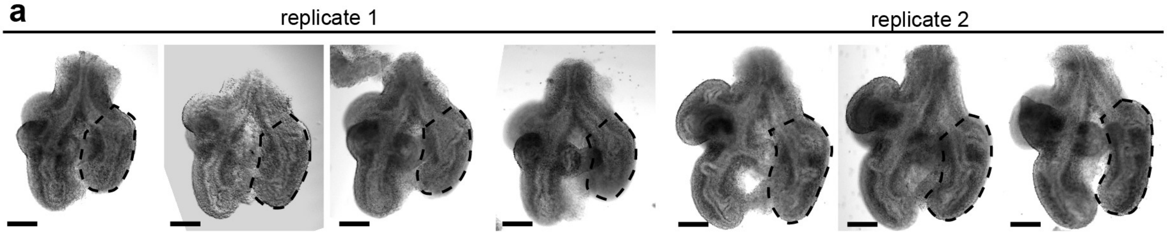


| b General | 1 | 2 |
|---|---|---|
| Estimated number of cells | 889 | 1082 |
| Mean reads per cell | 142396 | 119861 |
| Median genes per cell | 2779 | 1592 |
| Fraction of reads in cells | 71.50% | 65.10% |
| Total genes detected | 17529 | 17017 |
| Median UMI counts per cell | 9026 | 7713 |

| e Sequencing | 1 | 2 |
|---|---|---|
| Number of reads | 1E+08 | 1E+08 |
| Valid barcodes | 97.80% | 97.90% |
| Sequencing saturation | 78.10% | 80.30% |
| Q30 bases in barcode | 97.90% | 97.90% |
| Q30 bases in RNA read | 86.90% | 86.20% |
| Q30 bases in UMI | 98% | 98.10% |

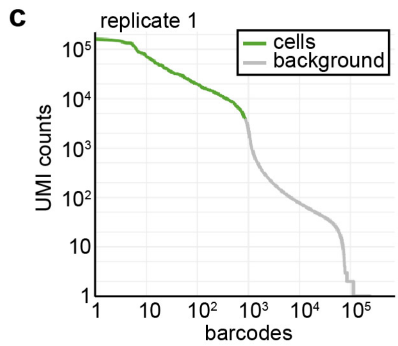


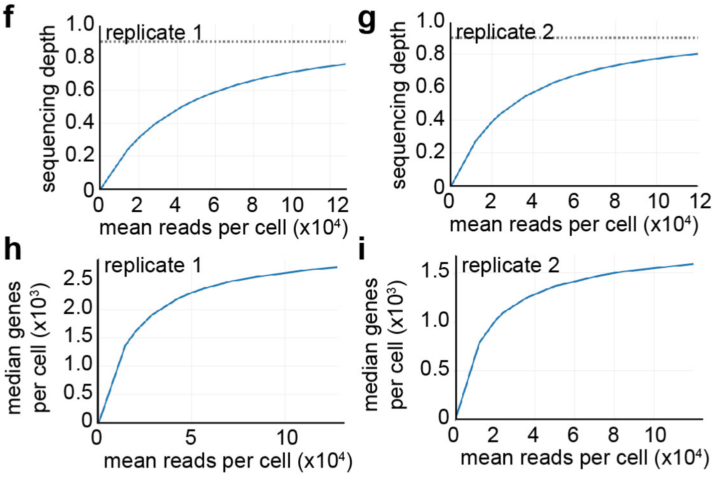


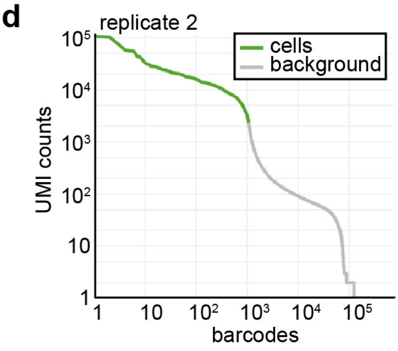


| j Mapping | 1 | 2 |
|---|---|---|
| Reads mapped confidently to genome | 73.30% | 72.10% |
| Reads mapped confidently to intergenic regions | 5.00% | 4.40% |
| Reads mapped confidently to intronic regions | 9.40% | 7.10% |
| Reads mapped confidently to exonic regions | 58.90% | 60.70% |
| Reads mapped confidently to transcriptome | 57.20% | 59.30% |
| Reads mapped antisense to gene | 0.80% | 0.60% |

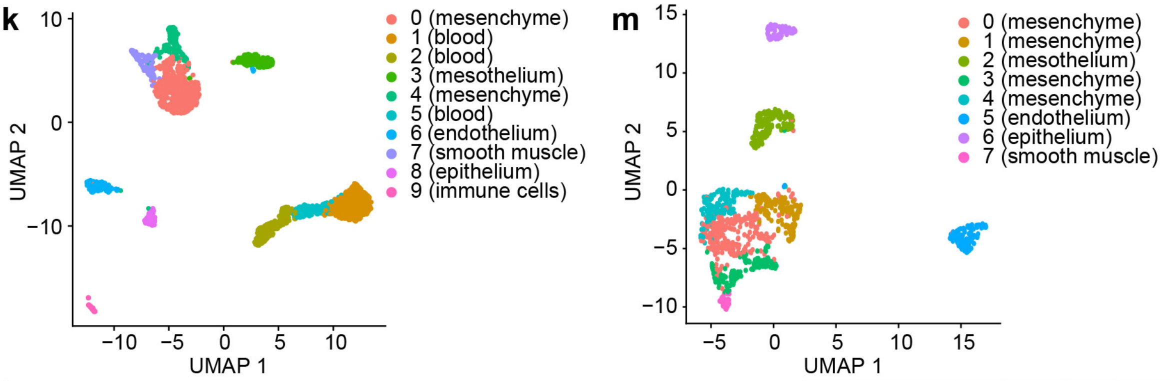


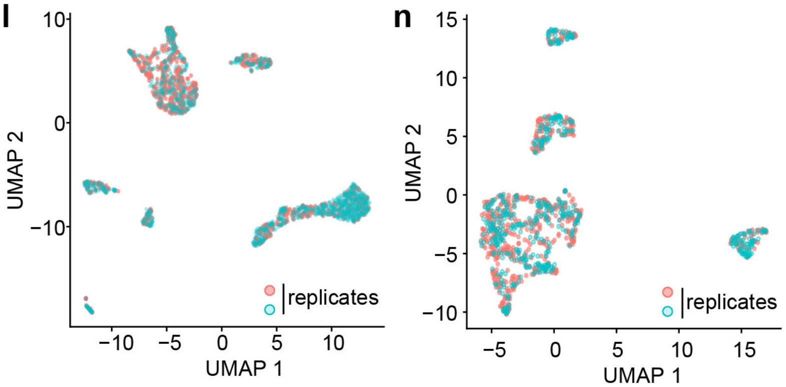


| o | before | | after | |
|---|---|---|---|---|
| number of clusters | 8 | | 6 | |
| | ID | # cells | ID | # cells |
| mesenchyme | 0 | 255 | 0 | 354 |
| | 1 | 163 | 1 | 328 |
| | 3 | 145 | | |
| | 4 | 138 | | |
| smooth muscle | 7 | 35 | 2 | 61 |
| epithelium | 6 | 79 | 3 | 79 |
| vascular endothelium | 5 | 121 | 4 | 117 |
| mesothelium | 2 | 155 | 5 | 152 |

### Supplementary Figure 2

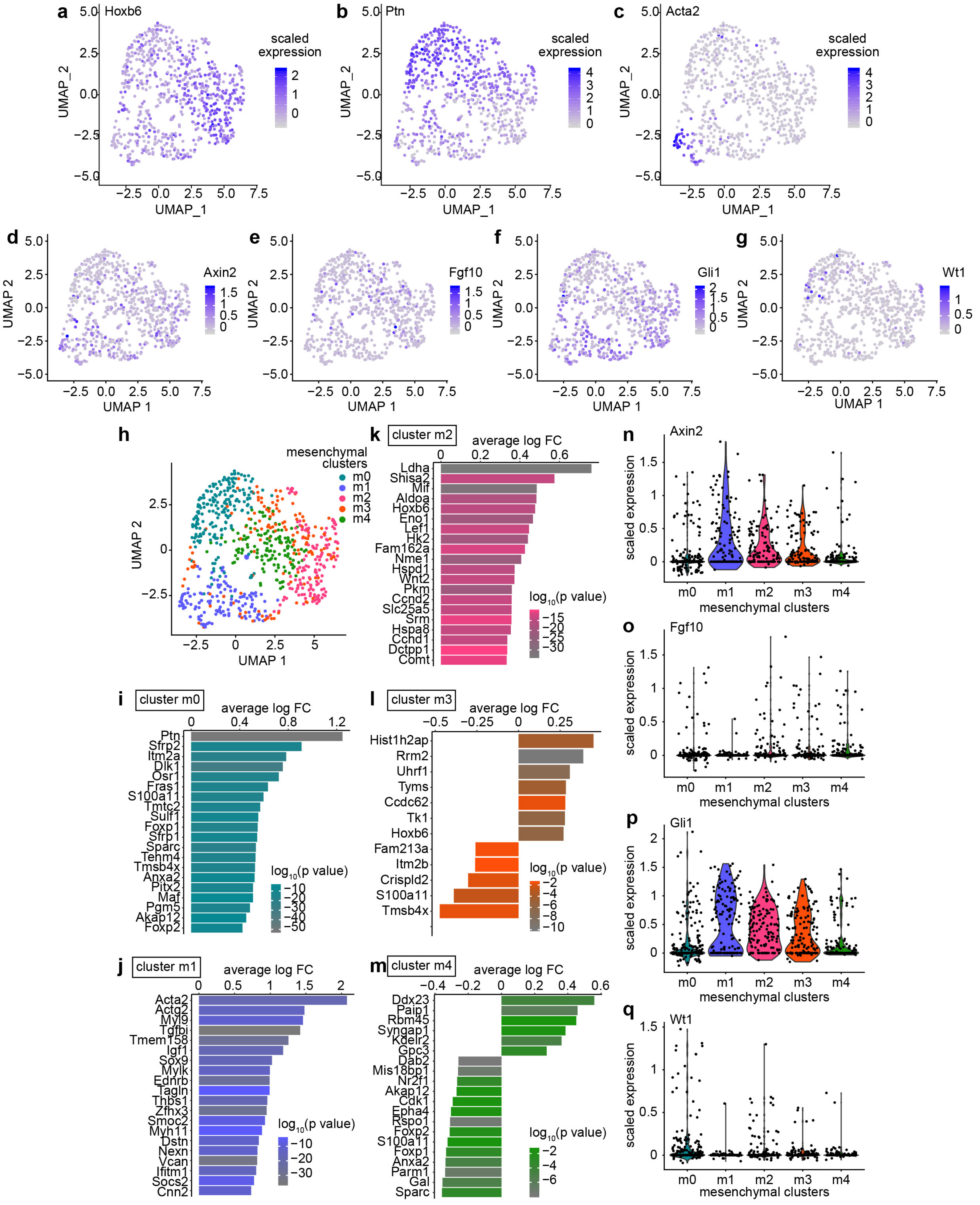

### Supplementary Figure 3

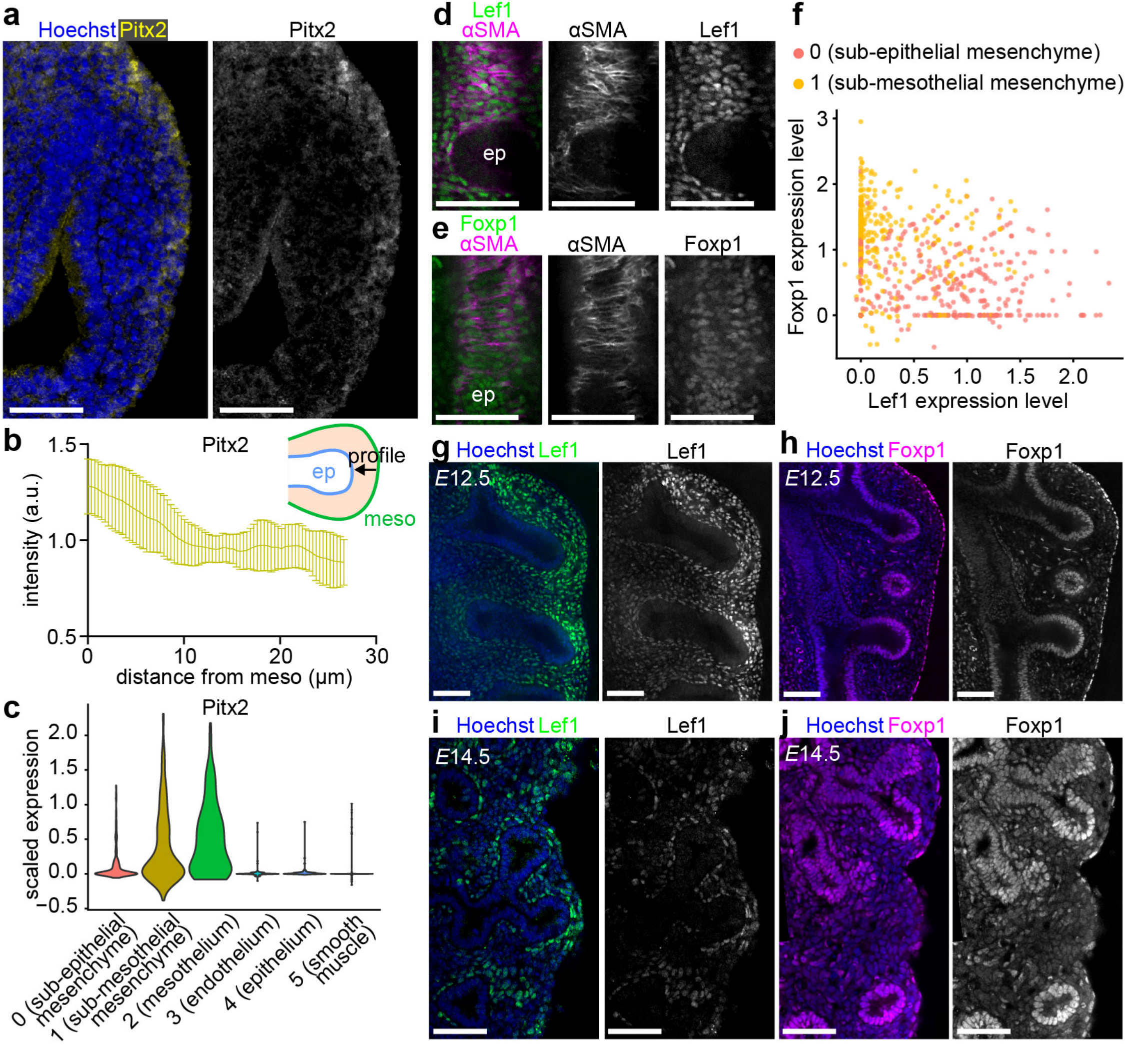

### Supplementary Figure 4

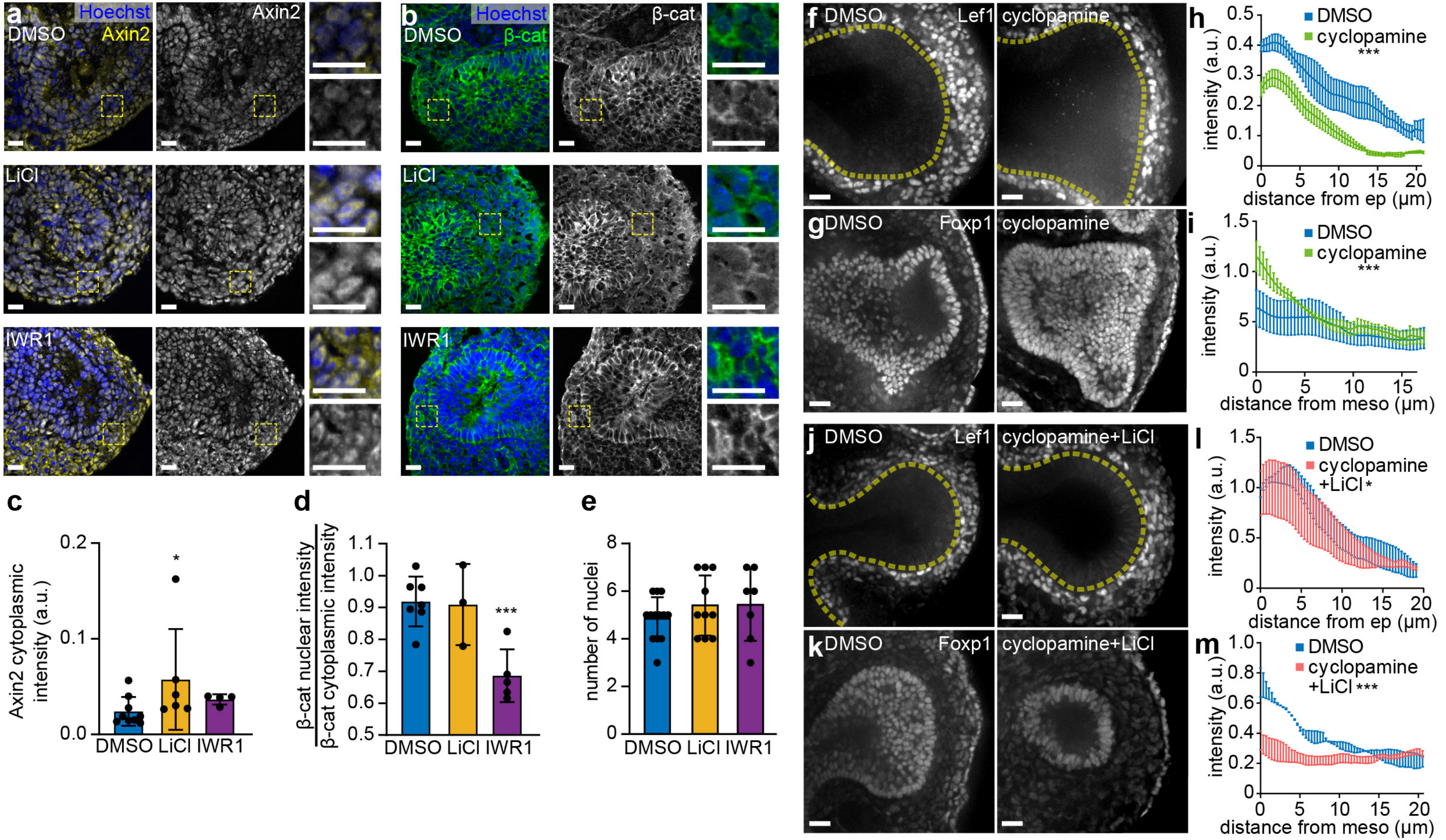

### Supplementary Figure 5

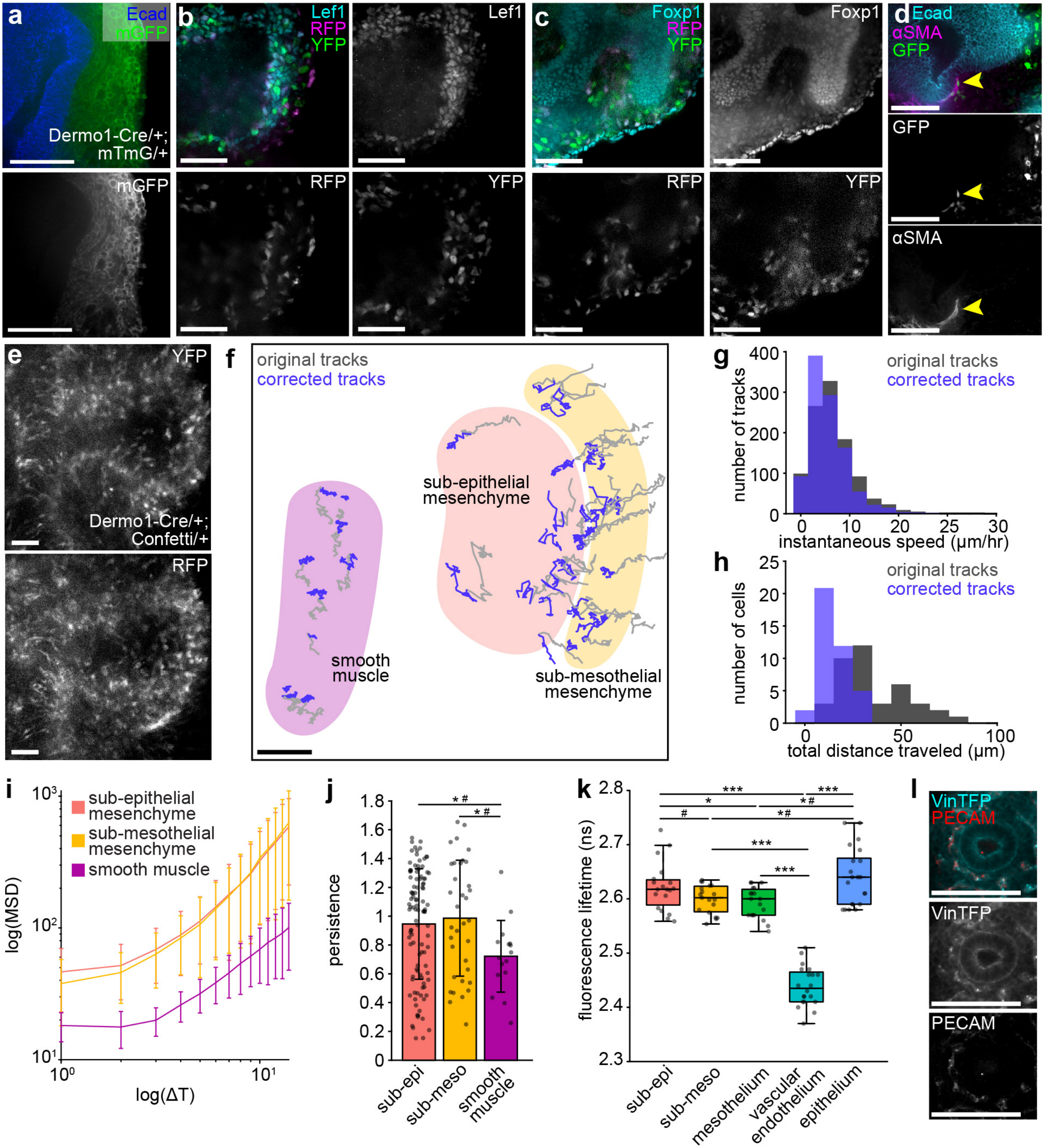

### Supplementary Figure 6

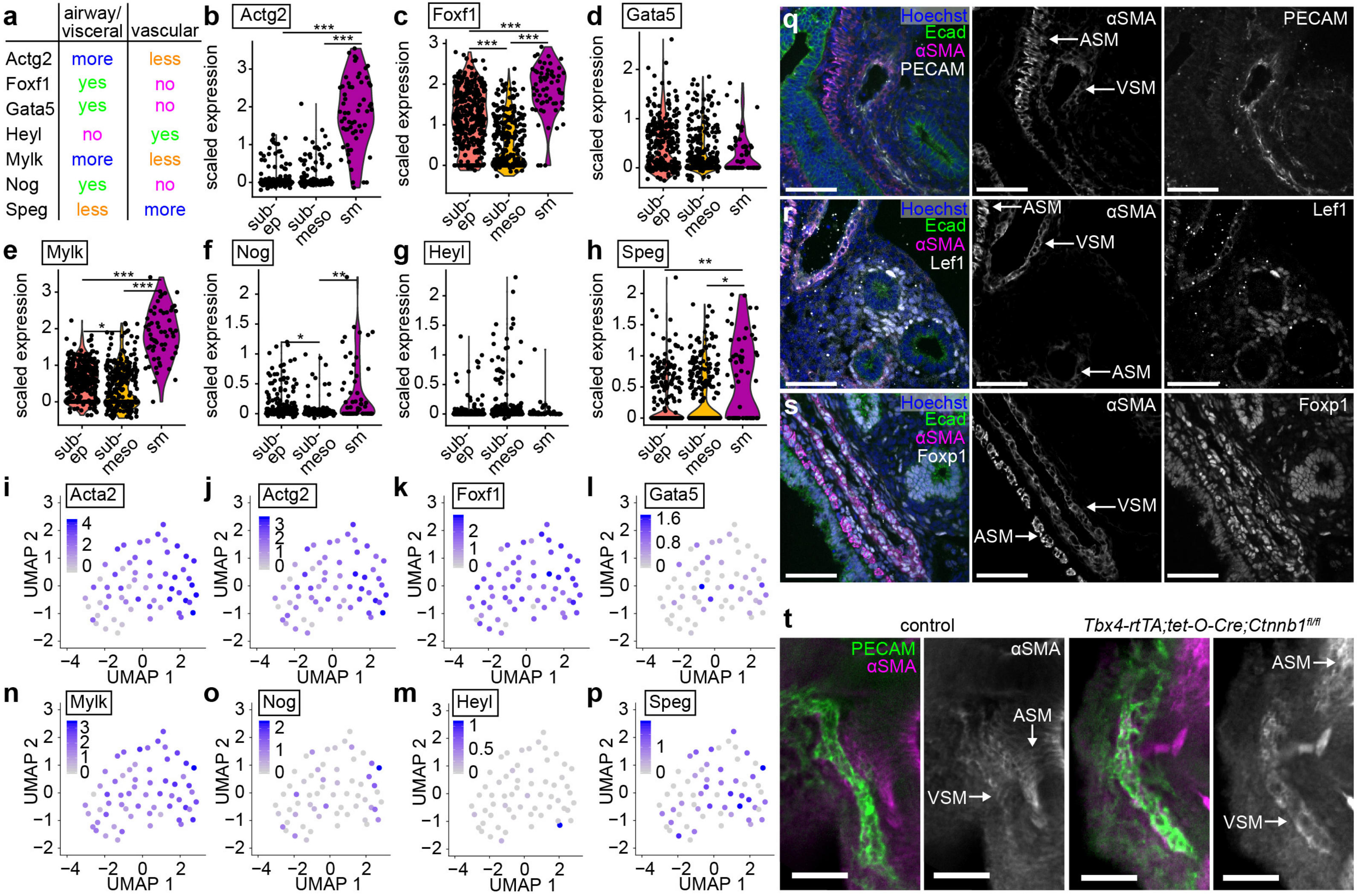

### Supplementary Figure 7

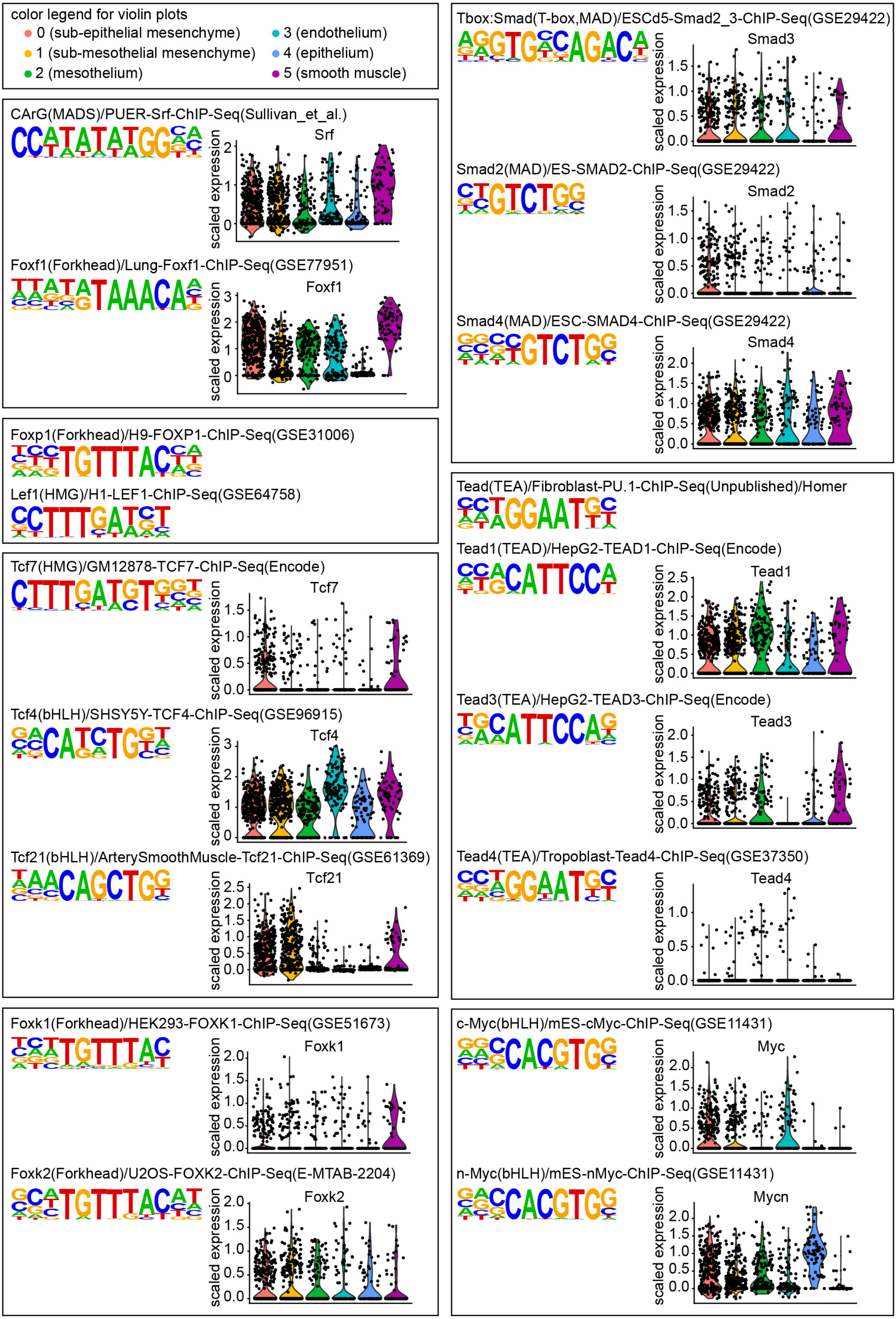

### Supplementary Figure 8

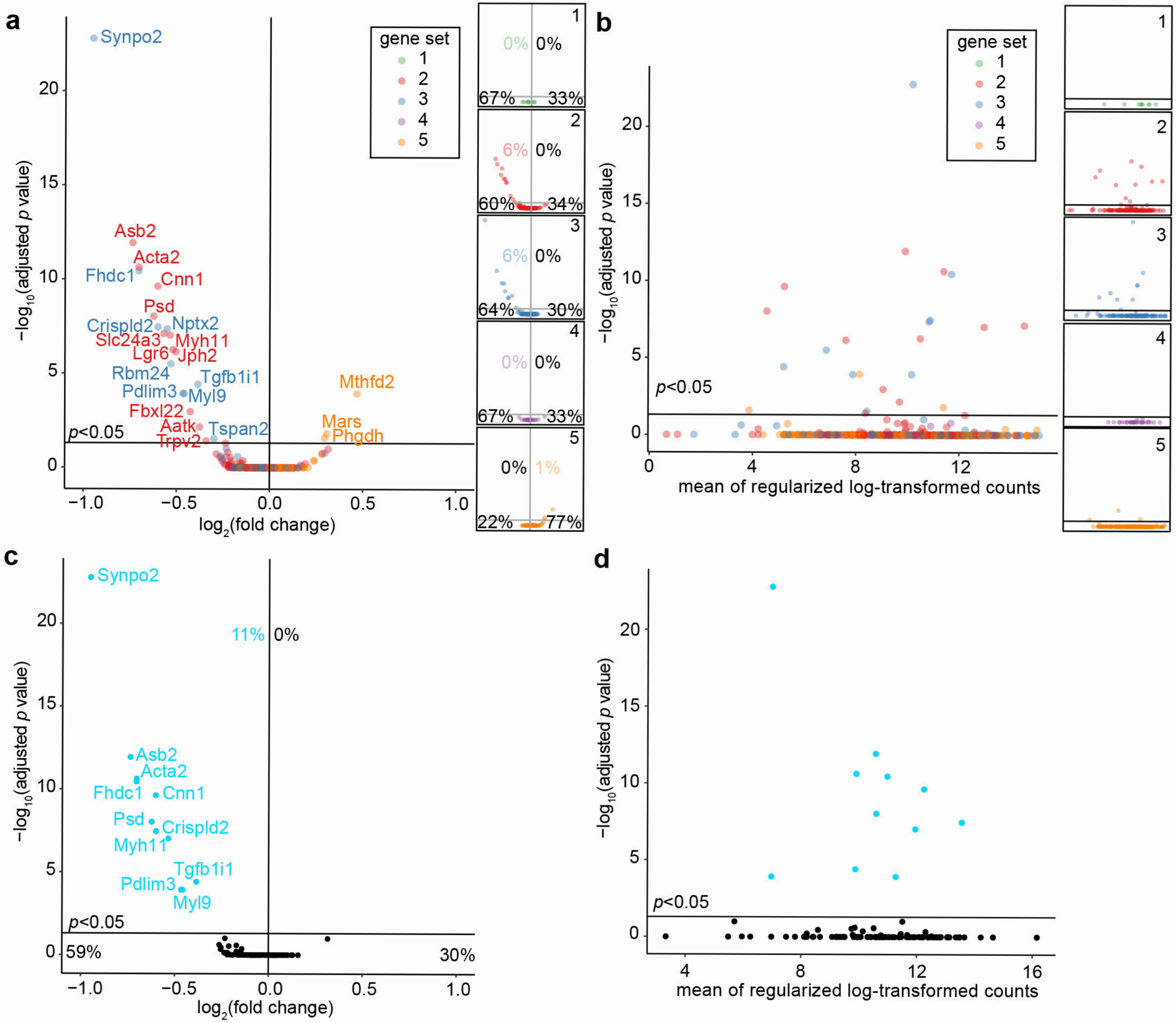

### Supplementary Figure 9

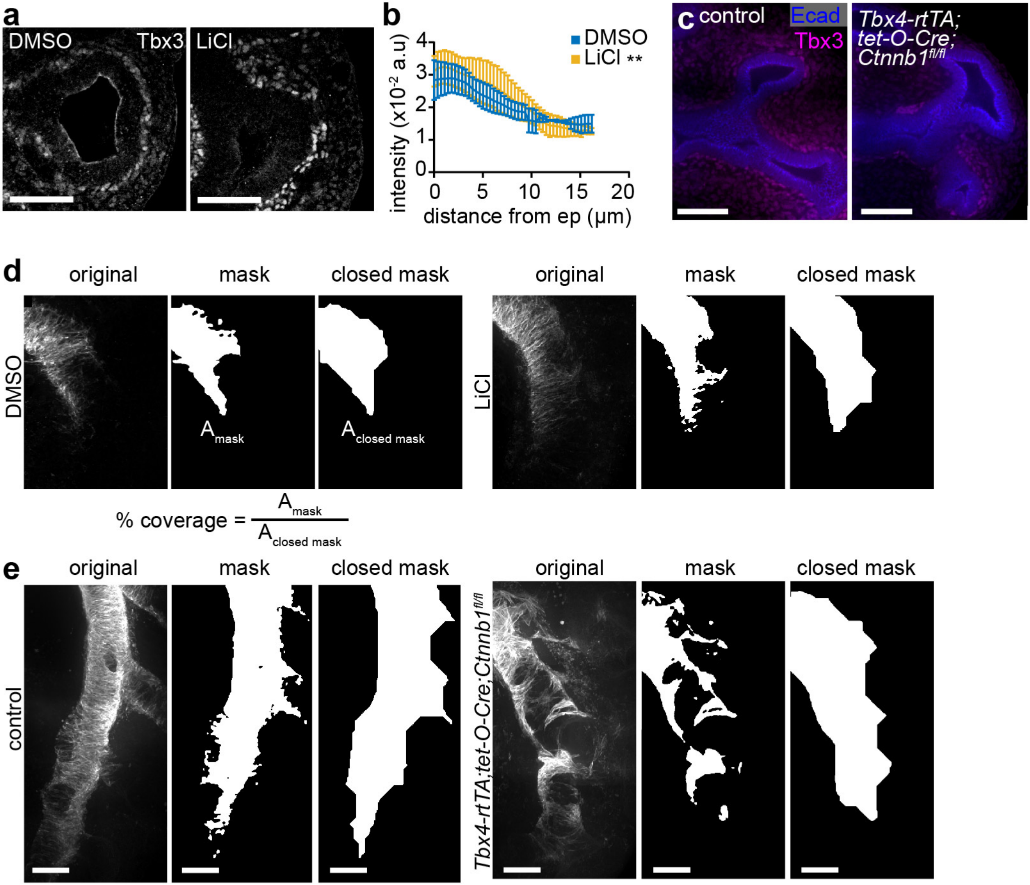
